# Functional and spatial proteomics profiling reveals intra- and intercellular signaling crosstalk in colorectal cancer

**DOI:** 10.1101/2022.09.16.508204

**Authors:** Christina Plattner, Giorgia Lamberti, Peter Blattmann, Alexander Kirchmair, Dietmar Rieder, Zuzana Loncova, Gregor Sturm, Stefan Scheidl, Marieke Ijsselsteijn, Georgios Fotakis, Asma Noureen, Rebecca Lisandrelli, Nina Böck, Niloofar Nemati, Anne Krogsdam, Sophia Daum, Francesca Finotello, Antonios Somarakis, Alexander Schäfer, Doris Wilflingseder, Miguel Gonzalez Acera, Dietmar Öfner, Lukas A. Huber, Hans Clevers, Christoph Becker, Henner F. Farin, Florian R. Greten, Ruedi Abersold, Noel Filipe da Cunha Carvalho de Miranda, Zlatko Trajanoski

**Affiliations:** Biocenter, Institute of Bioinformatics, Medical University of Innsbruck, Austria; Department of Biology, Institute of Molecular Systems Biology, ETH Zurich, Switzerland; Department of Visceral, Transplant and Thoracic Surgery, Medical University of Innsbruck, Austria; Department of Pathology, Leiden University Medical Center, Leiden, The Netherlands; Department of Radiology, Leiden University Medical Center, Leiden, The Netherlands; Institute of Hygiene and Medical Microbiology, Medical University of Innsbruck, Austria; Department of Medicine 1, University of Erlangen-Nuremberg, Erlangen, Germany; Biocenter, Institute of Cell Biology, Medical University of Innsbruck, Austria; Hubrecht Institute, Utrecht, The Netherlands; Institute for Tumor Biology and Experimental Therapy, Georg-Speyer-Haus Frankfurt/Main, Germany; Frankfurt Cancer Institute, Goethe University, Frankfurt am Main, Germany; German Cancer Consortium (DKTK), Heidelberg, Germany

**Keywords:** Kinase networks, functional precision medicine, patient-derived organoids, phosphoproteomics, imaging mass-cytometry

## Abstract

**Background:** Despite major advances in the development of targeted therapies, precision (immuno)oncology approaches for patients with colorectal cancer continue to lag behind other solid cancers. Functional precision oncology – a strategy that is based on perturbing primary tumor cells from cancer patients with drugs – could provide an alternate road forward to personalize treatment.

**Methods:** We extend here the functional precision oncology paradigm to measuring phosphoproteome landscapes using patient-derived organoids (PDOs). We first employed steady-state multi-omics (exome sequencing, RNA sequencing, and proteomics) and single-cell characterization of the PDOs. The PDOs were then perturbed with kinase inhibitors (MEKi, PI3Ki, mTORi, TBKi, BRAFi, and TAKi), and large-scale phosphoproteomics profiling using data-independent acquisition was carried out. Further, we used imaging mass-cytometry-based single-cell proteomic profiling of the primary tumors to characterize cellular composition of the tumor-microenvironment (TME) and to quantify heterocellular signaling crosstalk.

**Results:** We show that kinase inhibitors induce profound off-target effects resulting in a crosstalk with oncogenic and immune-related pathways. Reconstruction of the topologies of the kinase networks revealed that the patient-specific rewiring of the central EGFR-RAS-MAPK network is unaffected by mutations. Moreover, we show non-genetic heterogeneity of the PDOs and patient- and inhibitor-specific upregulation of stemness and differentiation genes by kinase inhibitors. We complemented our functional profiling by spatial proteomics profiling of the primary tumors using imaging mass cytometry. We quantify spatial heterocellular crosstalk and tumor-immune cell interactions, showing an avoidance of PD1+ immune cells and PD-L1+ tumor cells.

**Conclusions:** Collectively, we provide a multi-modal framework for inferring tumor cell intrinsic signaling and external signaling from the TME to inform precision (immuno)-oncology in colorectal cancer.

## Background

In the past two decades, tremendous advances have been made in both cancer biology by identifying recurrent mutations in oncogenic signaling pathways using sequencing technologies and therapeutics by developing targeted drugs specific for these mutations. In colorectal cancer (CRC), one of the major cancers with high incidence and where mortality rates are still high [1], progress in targeted therapy has been limited, relative to other solid cancers like lung cancer or melanoma [2]. The genetic heterogeneity [3] as well as paucity of druggable targets (nearly 50% of all CRCs are driven by un-druggable oncogenes of the *RAS* family) [2] pose considerable challenges for developing precision oncology approaches for patients with CRC. Moreover, CRC seems to be refractory to therapy with immune checkpoint blockers (ICB) with the notable exception of CRC tumors characterized by mis-match-repair deficiency or POLE proofreading mutations [4]. This is somehow paradoxical since CRCs, irrespectively of mismatch repair status, are known to be under immunological control, as we have shown in the past [5].

In precision oncology the paradigm is emerging that genomic profiling of the tumor assessed at early intervention (biopsy or resection) does not provide sufficient information to guide therapy. A diagnostic approach that is functional, i.e. measuring responses to perturbations with living cells derived from the specific tumor is expected to provide immediately translatable, personalized treatment information [6]. Such functional precision medicine approach requires patient derived models representing the tumors from affected individuals like patient-derived xenografts (PDX) or patient-derived organoids (PDOs) [7]. However, despite earlier encouraging reports [8,9], conflicting results have been reported regarding the capability of PDOs to predict tumor responses to specific drugs [10] Moreover, a recent study showed that despite sensitivity of the PDOs to selected kinase inhibitors, CRC patients from which these cells originated did not demonstrate objective clinical response to the treatment [11]. A number of reasons can be attributed to the limited value of PDOs as a tool for functional precision oncology. These include PDO culture success rate, ineligibility of the patients, or limited set of drugs tested [11]. While improvements for more streamlined experimental design are conceivable and could accelerate the implementation of PDOs to guide treatment decisions, major limitations of this drug-screening approach remain. The read-outs of the assays for testing *in vitro* response are based on growth rates of the cells and do not measure changes in the levels of functional status of the corre-sponding proteins and hence, do not provide insights into the mechanisms underlying sensitivity or resistance to a specific drug. Moreover, given the large number of available approved drugs, testing existing drugs for novel therapeutic strategies (drug repurposing) or testing novel combinations even for a limited number of single agents becomes impractical. Thus, an improved approach is needed that identifies key cancer cell vulnerabilities and provides rationale to select drugs/drug combinations. Given the fact that dysfunctional signaling in tumors arises from rewiring of signaling pathways and that nearly all molecularly targeted therapeutics are directed against signaling molecules [12], a strat-egy that focuses on cancer cell signaling measurements in PDOs could offer an alternate road forward.

Such a functional precision oncology strategy based on comprehensive dissection of tumor cell signaling depends on obtaining quantitatively accurate and consistent phosphoproteomics profiles. Recently, data-independent acquisition (DIA) methods emerged as a technology that combines deep proteome coverage with quantitative consistency and accuracy. Specifically, a variant of DIA methods called sequential window acquisition of all theoretical mass spectra (SWATH-MS)[13] was developed in which all ionized peptides within a specified mass range are fragmented for each sample in a systematic manner [14], and thereby enable reproducible high-throughput identification and quantification of proteomes across many samples. We reasoned that functional precision profiling using PDOs and SWATH-MS-based quantitative phosphoproteomics would enable patient-level reconstruction of kinase signaling networks and shed light on the intrinsic biology of the CRC cells.

We therefore first established a living biobank of PDOs from CRC patients and carried out steady-state proteogenomic characterization using DNA- and RNA-sequencing, and SWATH-MS-based proteomics (Figure 1). We then developed a functional precision oncology approach based on perturbation experiments of the PDOs with kinase inhibitors, quantitative phosphoproteomic measurements, and integration of *a priori* knowledge. We show that kinase inhibitors induce profound off-target effects that impact oncogenic and immune-related pathways. Reconstruction of the topologies of kinase signaling networks showed that patient-specific rewiring is largely unaffected by mutations. More- over, we show non-genetic heterogeneity of the PDOs and upregulation of stemness and differentiation genes by kinase inhibitors. Finally, we complemented our functional precision profiling by imaging mass cytometry (IMC)-based analysis of the primary tumors that enabled us to quantify spatial heterocellular crosstalk and tumor-immune cell interactions.

**Figure 1.**
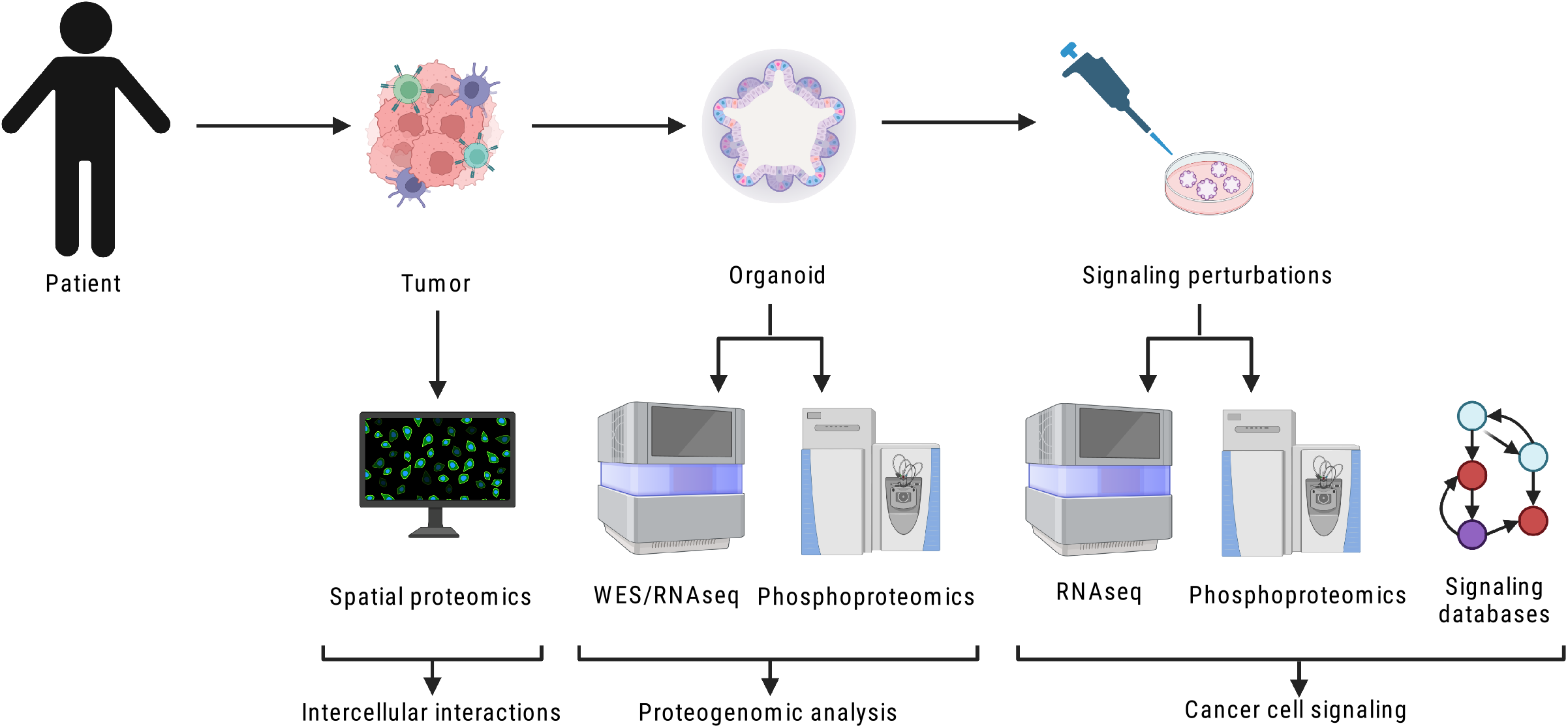
Schematic outline of the overall concept used in this study. Multimodal profiling of tumor specimens and PDOs in a cohort of CRC patients.

## Methods

### Human specimens

Histologically verified primary and metastatic colorectal tumor tissues and blood samples were obtained from patients undergoing surgical resection at the Medical University Hospital of Innsbruck. Samples were obtained from adult female or male patients who were treatment-naïve, with the exception of patient CRC26 who received FOLFOX and cetuximab before surgery. Written informed consent for research was obtained from patients prior to tissue acquisition. The medical ethical committee of the Medical University of Innsbruck approved protocol AN2016-0194 366/4.9 for the establishment of colorectal cancer PDOs cultures.

### Tumor cells isolation

Tumors were washed three times with DPBS (Thermo Scientific, Cat#14190169) containing 100 μg/ml Primocin (Invivogen, Cat#ant-pm-2) and 10 ml/L Penicillin-Streptomycin (Sigma, Cat#P4333), minced finely and incubated with 25 ng/ml Liberase (Roche, Cat#5401054001) in StemPro hESC SFM (Thermo Scientific, Cat#A1000701) for 1 hour at 37°C. After incubation, StemPro hESC SFM containing 10% FBS (Sigma, Cat#F7524) was added and the mixture was put over a 400 μM and a 100 μM cell strainers (pluriSelect, # 43-50100-51 and 43-50400-03) to remove large fragments. Cells were spun at 1000 rpm for 4 minutes, pellet was resuspended in 1x RBC Lysis Buffer (Biolegend, Cat#B420301) and incubated for 10 minutes at room temperature. Cells were spun at 1500 rpm for 5 minutes and pellet was washed three times with DPBS followed by centrifugation at 1500 rpm for 3 minutes.

### PDOs culture

Isolated tumor cells were seeded at a density of 1.5×10^5^ in 30 μl droplets of 70% Geltrex (Thermo Scientific, Cat#A1413202). The composition of PDO culture medium was: Advanced DMEM/F12 (Thermo Scientific, Cat#12634028) supplemented with 10 mM Hepes solution (Sigma, Cat#H0887), 10 ml/L Penicillin-Streptomycin solution, 2 mM GlutaMAX (Thermo Scientific, Cat#3550061), 20% Rspondin conditioned medium, 10% Noggin conditioned medium, 20 ml/L B-27 supplement (Thermo Scientific, Cat#17504044), 1.25 mM N-Acetylcysteine (Sigma, Cat#A9165), 0.5 nM A83-01 (Tocris, Cat#2939), 10 μM SB202190 (Sigma, Cat#S7067), 50 ng/ml human EGF (Peprotech, Cat#AF-100-15), 100 μg/ml Primocin (Invivogen, Cat#ant-pm-2), and 10 μM Y27632 (AbMole, Cat#M1817). PDO culture medium was refreshed every two days. To passage the PDOs, Geltrex was broken with a cell scraper and PDOs were collected in a tube. The PDOs were centrifuged at 1500 rpm for 5 minutes, medium was removed, the pellet was resuspended in TripLE Express (Thermo Scientific, Cat#12604013) and incubated for 5 minutes at 37°C. Advanced DMEM/F12 was added and cells were spun down at 1500 rpm for 5 minutes. The pellet was resuspended in 70% Geltrex and plated in 30 μl droplets on 6 wells-plates (Sarstedt, #83.3920), 4 drops each well. After allowing Geltrex to solidify, PDO culture medium was added to the plates and PDOs were incubated at 37°C with 5% CO2.

### Perturbation experiments with PDOs

PDOs were cultured and expanded to forty-eight 30μl culture-droplets for each condition. At 2 hours before collection, PDOs were treated with single inhibitors (or DMSO as solvent control) or at 1 hour before collection were stimulated with ligand (or solvent control H2O). The following inhibitors and ligand were used as previously evaluated [15]: MEKi AZD6244 (4 μmol/L, Biomol, #LKT-S1846.1), PI3Ki AZD6482 (10 μmol/L, Biomol, #Cay15250-1), mTORi AZD8055 (2 μmol/L, Eubio, #SYN-1166-M001), TBK1i BX-795 (10 μmol/L, Sigma, #SML0694), BRAFi PLX4720 (5 μmol/L, Biomol, #Cay15142-1), TAK1i 5Z-7-Oxozeaenol (5 μmol/L, Sigma, O9890), and TNFα (10 ng/mL, Peprotech, AF-300-01). After treatment, cultivation dishes were placed on ice and PDOs culture-droplets were washed twice with ice-cold DPBS. Culture-droplets were disrupted in Cell Recovery Solution (Corning, #7340107), collected in a tube and incubated for 1 hour on ice. After incubation, PDOs were spun at 1500 rpm for 10 minutes at 4°C, washed twice with ice-cold DPBS and PDO pellets were snap-frozen in liquid nitrogen and stored at −80°C.

### PDOs preparation for proteomics analysis

PDOs were cultured and expanded to six 30μl culture-droplets for each sample. At the time of collection, cultivation dishes were placed on ice and PDOs culture-droplets were washed twice with ice-cold DPBS. Culture-droplets were disrupted in Cell Recovery Solution (Corning, #7340107), collected in a tube and incubated for 1 hour on ice. After incubation, PDOs were spun at 1500 rpm for 10 minutes at 4°C, washed twice with ice-cold DPBS and PDO pellets were snap-frozen in liquid nitrogen and stored at −80°C.

### DNA and RNA sequencing

PDOs were harvested, snap-frozen and their DNA were extracted using the PureLink Genomic DNA Mini Kit (Thermo Scientific, #K182001) following manufacturer instructions. Germline DNA were extracted from frozen peripheral blood mononuclear cells (PBMCs) using PureLink Genomic DNA Mini Kit (Thermo Scientific, #K182001) following manufacturer instructions. RNA was isolated from PDO pellets using the RNeasy Plus Mini Kit (Qiagen, #74134) following manufacturer instruction. Exome-sequencing was performed using SureSelect all human V6 capture kit and Illumina sequencing (GATC, Konstanz, Germany and GENEWIZ, Leipzig, Germany). Total RNA was isolated from frozen PDO pellets using the RNeasy Plus Mini Kit (Qiagen, #74134) following manufacturer’s instruction and submitted to total transcriptome full-length mRNA sequencing (GATC, Konstanz, Germany, Medical University of Innsbruck).

### Single cell sequencing

PDO culture-droplets were washed once with warm DPBS, disrupted with a cell scraper and collected in a tube followed by centrifugation at 1500rpm for 5 minutes at room temperature. PDO pellets were resuspended in Trypsin-EDTA (Sigma, Cat#T4174) using 500 μl for each culture-droplet used, resus-pended 5 times with a 1000μl tip and incubated for 5 minutes at 37°C. An equal volume of Advanced DMEM/F12 was added and PDOs were further dissociated mechanically by resuspending 10 times using a 200 μl pipette tip placed on top of a 1000 μl tip. Cells were filtered through a 40 μm cell strainer and centrifuged at 1500rpm for 5 minutes at 4°C. Supernatant was removed and cell pellets were resuspended in 1 ml ice-cold 0.04% BSA in DPBS. Cells were counted with a hemocytometer (Marienfeld Neubauer, Cat#0640010) and viability was assessed using Trypan-blue solution (Sigma, Cat#T8154). Single cell suspensions of freshly isolated cells were converted to indexed scRNAseq libraries, using the Chromium Single Cell 3’GEX V3.1 technology from 10x Genomics, aiming for 8.000 cells per library. The resulting Libraries were sequenced with Illumina Novaseq technology (sequencing performed at Genewiz, Leipzig, Germany).

### Immunofluorescence

PDO samples for immunofluorescence were prepared as described [16]. Briefly, PDOs were freed from Geltrex by incubation in Cell Recovery Solution (Corning, #7340107) for 1 hour on ice, fixed in 4% PFA (Sigma, #16005) in PBS for 1 hour at RT and permeabilized in 1% Triton X-100 (Sigma, #T9284) for 30 minutes at RT. PDOs were incubated in Blocking Buffer (10% goat serum (Sigma, #G9023), 0.2% Triton X-100, 5% BSA in PBS) for 1 hour at RT and with primary antibodies in Blocking Buffer overnight at 4°C. PDOs were washed twice with PBS, incubated with secondary anti-bodies in Blocking Buffer for 2 hours at RT in the dark, washed twice with PBS and mounted in Vec-tashield Antifade Mounting Medium with DAPI (#H-2000-2). Following primary antibodies were used: anti-chromogranin A (Santa Cruz Biotechnology, #sc-393941, 1:500), anti-lysozyme (Abcam, #ab108508, 1:500), anti-mucin 2 (Santa Cruz Biotechnology, #sc-515032, 1:500), anti-EpCAM (R&D, #AF960-SP, 1:20), anti-Ki67 (Abcam, #ab92742, 1:500), anti-Lgr5 (Abcam, #ab75732, 1:100), anti-SOX9 (Sigma, #AB55351:500), Phalloidin-iFluor 647 (Abcam, #ab176759, 1:1000). All secondary antibodies were used 1:800. Immunofluorescence images were captured with an Operetta CLS High-Content Analysis System (Perkin Elmer).

### Western blotting

PDOs were lysed in RIPA buffer with 1x EDTA-free protease inhibitor cocktail (Sigma, #11836170001), 1% v/v phosphatase inhibitor cocktail (Sigma, #P0044) and 2mM PMSF (Sigma, #93482). Protein concentration was measured by BCA Protein Assay Kit (Thermo Scientific, #23225), proteins were separated on 7%, 10% or 12% precast polyacrylamide gels (NuPAGE) following the manual. Blotting was performed using the Invitrogen™ Novex™ XCell™ SureLock™ Blot-Modul (Invitrogen) according to the manufacturer instructions. Following primary antibodies were used according to manufacturer’s protocol: GAPDH (Santa Cruz Biotechnology, #sc-47724, 1:1000 and Invitrogen, #AM4300, 1:10000), phospho-mTOR (Cell Signaling, #5536T, 1:2000), phospho-AKT (Cell Signaling, #4060S, 1:1000), phospho-ERK1/2 (Cell Signaling, #4377S, 1:1000), phosho-MEK1 (Invitrogen, #MA-32165, 1:1000), phospho-p38 MAPK (Invitrogen, #36-8500, 1:1000. Chemiluminescence was recorded using the Image Quant Las4000 imaging system (GE Healthcare).

### mRNA expression analysis of selected genes by RT-qPCR

PDOs were cultured in 6 well flat bottom plates and treated with kinase inhibitors (5 μM BRAFi, 4μM MEKi, 2μM mTORi, 10μM PI3Ki, 5μM TAKi, 0.25 μM TBKi) and vehicle control (DMSO) for 72 hrs. After the treatment, organoid pellets were harvested by washing twice in cold PBS, snap frozen in liquid N2 and stored at −80°C before RNA extraction. Total RNA was isolated using RNeasy Plus Mini Kit (Qiagen, cat#: 74134). cDNA was synthesized using SuperScriptIII first-strand synthesis system for RT-PCR (Invitrogen) with 1μg of total RNA as a template. Quantitative PCR (qPCR) was performed in MicroAmp™ Optical 384-Well Reaction Plates with Barcode (Applied Biosystems, cat#: 4309849) on ViiA™ 7 Real-Time PCR System (Applied Biosystems) using Platinum™ SYBR™ Green qPCR SuperMix-UDG w/ROX (Invitrogen, cat#: 11744500). The final volume for qPCR reaction was 6μl containing 4ng of cDNA, 0.2μM of each primer and 1X Platinum™ SYBR™ Green qPCR SuperMix. Cycling conditions for qPCR were as follows: 2 mins at 50°C, 10 mins at 95°C followed by 40 cycles of 15 s at 95°C and 60 s at 60°C. Primers used for qPCR are listed in Table S1.

### Sample preparation for mass spectrometric analyses

Pelleted and frozen PDOs were lysed in 8 M Urea (Sigma, #33247) in 100 mM ammonium bicarbonate (Sigma, #09830) and with sonication for 10 min. For the perturbation experiments, 1:100 phosphatase inhibitor cocktails (Thermo Scientific #P5726 and Sigma #P0044) were added to the lysis buffer. To reduce and alkylate the disulfide bonds, the lysate was reduced using 5 mM tris(2-carboxyethyl)phosphine (TCEP) for 30 min at 37°C and alkylated using 40mM Iodacetamide (IAA) for 45 min at 25°C in the dark. The protein amount was measured using the Bicinchoninic acid (BCA) assay (Thermo Scientific, #23225) and the appropriate protein amount (60 μg for baseline expression experiments and 1mg for perturbation experiments) was digested with LysC (1:100, Wako, #121-05063) for 4 h and Trypsin (1:75, Promega, #V5113) overnight. For the digestion with LysC and Trypsin, the samples were diluted to 6 M and 1.5 M Urea (Sigma, #33247) in 100mM ammonium bicarbonate respectively using 100 mM ammonium bicarbonate (Sigma, #09830). The digestion was stopped the following day by acidification with trifluoroacetic acid to pH 2-3.

For the baseline expression experiments, the digested peptides were desalted using NEST C18 Micro-Spin™ columns by washing with 2% acetonitrile 0.1% trifluoroacetic acid and eluting with 50% acetonitrile 0.1% trifluoroacetic acid. The eluted peptides were dried in a vacuum concentrator, reconstituted in 60μl 2% acetonitrile 0.1% formic acid in H2O, and spiked with iRT peptides (Biognosys, #Ki-3002-1) prior to injection into the mass spectrometer.

For the perturbation experiments, the digested peptides were desalted using Waters Sep-pak C18 columns by washing with 0.1% trifluoroacetic acid in H2O and eluting with 50% acetonitrile and 0.1% trifluoroacetic acid in H2O. The eluted peptides were subsequently dried in a vacuum concentrator. Before drying fully, an aliquot (1:20) was taken for the corresponding total cell lysate samples. The aliquot was dried and the peptides were dissolved in 2% acetonitrile and 0.1% formic acid in H2O and spiked with iRT peptides (Biognosys, #Ki-3002-1) prior to injection into a mass spectrometer. The remaining part of the sample was destined for phosphoenrichment and dried in a vacuum concentrator. To enrich for phosphopeptides, the peptides were first dissolved in a tube with loading buffer (1 M glycolic acid, 5% trifluoroacetic acid, 80% acetonitrile in H2O) by shaking 10 min and sonicating for 10 min. For phosphoenrichment, stage tips were constructed placing two C8 plugs into an empty 300 μl tip. These stage tips were placed in a tube using a connector and centrifuged on a table top centri-fuge at around 800 g for all subsequent washes. The stage tips were washed with 200 μl methanol to condition the filter. To the washed tips, 80 μl TiO2 bead slurry (2.5 mg TiO2 beads in 50% acetoni-trile, 0.1% trifluoroacetic acid) were added and the beads were equilibrated with 200 μl loading buffer. The peptides were loaded by starting with a low centrifugation force of 100 g that was progressively increased until all peptides were loaded. The loaded peptides were washed once with 100 μl loading buffer, once with 100 μl 80% acetonitrile and 0.1% trifluoroacetic acid in H2O, and once with 100ul 50% acetonitrile and 0.1% trifluoroacetic acid in H2O. The peptides were eluted with 250 μl 0.3 M NH3OH and a subsequent elution of 20 μl 50% acetonitrile and 0.1% trifluoroacetic acid in H2O to elute peptides from the filter paper. The phosphopeptides were eluted directly in a tube with trifluoroacetic acid to reach pH 2. The phosphopeptides were then desalted using NEST UltraMicro-SpinTM C18 columns and eluted with 50% acetonitrile, 0.1% trifluoroacetic acid in H2O. The buffer was evaporated in a vacuum concentrator and the peptides were dissolved in 2% acetonitrile and 0.1 % formic acid in H2O and spiked with iRT peptides (Biognosys, #Ki-3002-1) prior to injection of samples into a mass spectrometer.

### Acquisition of samples using mass spectrometry

For the perturbation experiments, the peptides were analyzed on an Orbitrap Fusion Lumos mass spectrometer (Thermo Scientific, San Jose, CA) connected to an Easy-nLC 1200 (Thermo Scientific, San Jose, CA) HPLC system. Between 1 μl and 4 μl of peptide solution was separated by nano-flow liquid chromatography using a 120 min gradient from 5 to 37% buffer B in buffer A (Buffer A: 2% acetonitrile, 98% H2O, 0.1% formic acid; Buffer B: 80% acetonitrile, 20% H2O, 0.1% formic acid) on an Acclaim PepMap RSLC 75 μm x 25cm column packed with C18 particles (2 μm, 100 Å) (Thermo Scientific, San Jose, CA). The peptides were ionized using a stainless steel nano-bore emitter (#ES542; Thermo Scientific) using 2000 V in positive ion mode.

To build the spectral libraries, the samples were acquired in data-dependent acquisition (DDA) mode. The DDA method consisted of a precursor scan followed by product ion scans using a 3 s cycle time. The precursor scan was an Orbitrap full MS scan (120’000 resolution, 2 × 105 AGC target, 100 ms maximum injection, 350-1500 m/z, profile mode). The product ion scans were performed using Quadrupole isolation and HCD activation using 27% HCD Collision Energy. The Orbitrap was used at 30’000 resolution and setting the RF Lens at 40%. The AGC Target was set to 5 × 105 and 50 ms maximum injection time. Charge states of 2-5 were targeted and the dynamic exclusion duration was 30s.

To quantify the peptide abundance, the samples were acquired in data-independent acquisition (DIA) mode. The DIA method consisted of a precursor scan followed by product ion scans using 40 windows between 400 m/z and 1000 m/z. The precursor scan was an Orbitrap full MS scan (120,000 resolution, 2 × 105 AGC target, 100 ms maximum injection, 350-1500 m/z, profile mode). The product ion scans were performed using Quadrupole isolation and HCD activation using 27% HCD Collision Energy. The Orbitrap was used at 30,000 resolution using a scan range between 200 and 1800 and setting the RF Lens at 40%. The AGC Target was set to 5 × 105 and 50 ms maximum injection time.

For the baseline expression experiments, the peptides were measured on a Sciex TripleTOF 5600 mass spectrometer with a 90 min gradient and the mass spectrometer was operated in SWATH mode. The precursor peptide ions were accumulated for 250 ms in 64 overlapping variable windows within an m/z range from 400 to 1200. Fragmentation of the precursor peptides was achieved by Collision Induced Dissociation (CID) with rolling collision energy for peptides with charge 2+ adding a spread of 15eV. The MS2 spectra were acquired in high-sensitivity mode with an accumulation time of 50 ms per isolation window resulting in a cycle time of 3.5 s. The samples from the different PDOs were injected consecutively in a block design to prevent any possible confounding effects due to deviation in machine performance. CRC03 and CRC26 were acquired at a later time point, but some original samples were reinjected in parallel to assess that the performance of the machine was similar.

### Building the spectral library for the perturbation experiments

The raw spectra were analyzed using MaxQuant version 1.5.2.8 that matched each spectrum against a FASTA file containing 20,386 reviewed human (downloaded on August 13, 2018 from www.uniprot.org) and iRT peptides and enzyme sequences. Carbamidomethyl was defined as a fixed modification, and Oxidation (M) as variable modifications. Standard MaxQuant settings for Orbitrap were used (e.g. peptide tolerance 20 ppm for first search and 4.5 for main search). In total, two searches were performed involving 54 injections of peptides and they resulted in the identification of 42’424, peptides from 4’239 protein groups, respectively. The four searches were imported into Spectronaut Pulsar (version 14.0.200309.43655 (Copernicus) Biognosys, Schlieren) to build spectral libraries with the following settings: PSM FDR Cut off of 0.01, fragment m/z between 200 and 1’800, peptides with at least 3 amino acids, fragment ions with a relative intensity of at least 5, precursors with at least 5 fragments. Moreover, an iRT calibration was performed with a minimum root mean square error of 0.8 and segmented regression. The spectral library for the total cell lysates contained coordinates for 54’551 precursor peptides from 4’223 protein groups. The spectral library for the phosphopeptide contained coordinates for 30’969 precursor peptides from 4’605 protein groups.

### Extraction of quantitative data from the mass spectrometry spectra

For the perturbation experiments, quantitative data were extracted from the acquired SWATH-MS maps using Spectronaut Pulsar (version 14.0.200309.43655 (Copernicus) Biognosys, Schlieren) (version 14). As SWATH Spectral library, we used our in-house compiled spectral libraries for the PDOs (see above). We used standard settings (they include a dynamic MS1 and MS2 mass tolerance strategy, a dynamic XIC RT Extraction Window with a non-linear iRT calibration strategy, and identification was performed using a precursor and protein Q value cutoff of 0.01). The quantified intensities for each fragment were extracted from 104 (phospho-enriched samples), 99 (total cell lysate) SWATH-MS injections and the fragment intensities were exported for further statistical analysis to R. Only quantities for fragments that have been detected at least two times in a given condition were selected. Further filtering was performed with mapDIA where a standard deviation factor of 2 and a minimal correlation of 0.25 were used to filter for reliable fragments.

For the baseline expression experiments, the SWATH-MS data was quantified using the OpenSWATH workflow on the in-house iPortal platform using the PanHuman Library [17]. An m/z fragment ion extraction window of 0.05 Th, an extraction window of 600 s, and a set of 10 different scores were used as described before. To match features between runs, detected features were aligned using a spline regression with a target assay FDR of 0.01. The aligned peaks were allowed to be within 3 standard deviations or 60 s after retention time alignment. The data was then further processed using the R/Bioconductor package SWATH2stats.

### Variant calling, Copy Number Variation and Neoantigen prediction

Somatic mutations, copy number alterations, Class I and II HLA types and neoantigens were called by running our previously published neoantigen prediction pipeline nextNEOpi [18] (version 1.1).

Briefly, we used the pipelines default options but enabled automatic read trimming to remove adapter sequence contamination from raw WES and RNAseq reads and we disabled NetChop and NetMHCstab. Further, we created a panel of normals from the healthy PBMCs and used it to identify recurrent technical artefacts in order to improve the results of the variant calling analysis. Finally, predicted neoantigens were filtered and prioritized using the “relaxed” filter set from nextNEOpi. The MSI status was determined with MSIsensor [19] and the scores were plotted as bar plots.

### RNA-sequencing data analysis

Sequence reads were preprocessed and mapped to the human genome GRCh38/hg38 and GENCODE v33 annotations using the nf-core RNAseq pipeline version 1.4.2 (git revision ff4759e)[20]. In brief, reads were mapped using STAR v2.7.1a [21] and gene expression quantified with RSEM v1.3.3 [22]. Pathway activity scores were estimated from normalized raw counts using the PROGENy (Pathway RespOnsive GENes) method [23]. GO enrichment analysis was performed using the R package ClusterProfiler 4.1.0 [24]. Eight major clusters were defined and annotated by the GO enrichment results. Heatmaps for visualization of RNAseq results were generated using the ComplexHeatmap 2.9.0 R package [25].

### Single-cell RNA-sequencing data analysis

The single-cell RNA sequencing reads were mapped to GRCh38-2020-A reference provided by 10x Genomics using the CellRanger pipeline (v5.0.0). Cellranger’s pre-filtered count matrices were loaded into scanpy [26] and filtered based on the following quality control cutoffs: ≥ 2000 genes, 2000 ≤ counts ≤ 75,000, < 25% mitochondrial reads. Doublets were removed using SOLO [27] v0.6.0. The 5000 most highly variable genes (HGVs) were detected with *scanpy*.*highly_variable_genes* with *flavor=“seurat_v3”* and *batch_key=“organoid”*. Batch effects were removed using scvi-tools v0.11.0 [28] based on the HVGs and using PDOs as the batch key. A neighborhood graph and UMAP embedding [29] were calculated with scanpy based on the SCVI latent representation and otherwise default parameters. Unsupervised clustering with the leiden-algorithm [30] based on the SCVI-corrected neighborhood graph (resolution=0.5) yielded 12 clusters. Marker genes for each cluster were detected using scvi-tools differential gene expression module [31] and clusters manually assigned to 7 epithelial cell-types based on these marker genes. For visualization, gene expression was CPM-normalized and log1p-transformed, before computing an additional (uncorrected) neighborhood graph and UMAP embedding. RNA velocity was estimated using velocyto.py (v0.17.17) [32] based on cellranger outputs using the *run10x* command and subsequently loaded into scvelo [33] for visualization. We performed single-cell pathway analysis using PROGENy [23,34]. Scores were computed using the *progeny-py* package v1.0.6. The top 1,000 target genes of the progeny model were used, as recommended for single-cell data.

### Proteomic data analysis

After quality control steps the median intensity value of baseline proteomic triplicates was retrieved for downstream analysis. Non-uniquely identified proteins were also discarded from further analyses. Differential abundance analysis was done using mapDIA version 3.1.0 [35]. Single-sample enrichment analysis of baseline protein expression levels was performed with the GSVA R package version 1.42.0 [36] using HALLMARK gene sets (version 7.5.1) imported with msigdbr 7.5.1 from MSigDB [37]. All samples, including replicates were transformed to log2 scale and resulting enrichment scores were then averaged per organoid. GO enrichment analysis was performed the same way as for the RNAseq data (8 major clusters). Heatmaps for visualization of proteomics results were generated using the ComplexHeatmap 2.9.0 R package [25].

### Correlation between mRNA and protein abundance

In total, the dataset comprises 3723 overlapping genes and proteins, but only those which were present in at least four out of the 12 matching samples were considered for correlation analysis (n=3536). Log2 transformed TPM-values for mRNA and log2-transformed protein abundances were used to calculate the Pearson correlation coefficient for each gene. The average correlation between mRNA and protein abundance is 0.29. The results were visualized in a histogram using the ggplot2 R package.

### Protein complexes

The list of manually curated protein complexes was retrieved from Ori et al [38]. Human protein complexes with a minimum of five proteins were selected and Pearson correlations within complexes were calculated across PDOs. The top 25% of the protein complexes were considered as variable, whereas the bottom 25% were defined as stable protein complexes. For the 26S proteasome the variance was calculated for all members of the complex, ignoring missing values. The results were visualized as a barplot using the ggplot2 R package.

### Phosphoproteomic data analysis

Phosphopeptide fragment data were prefiltered for intensities above 2000 and peptides with at least five measured fragments. Missing replicate values were replaced by the 20% value of the minimum of the corresponding fragment abundance. Differential abundance analysis was done using mapDIA version 3.1.0 [35]. Protein sequences and gene symbols were retrieved from UniProt [39] using the UniProt.ws 2.36.5 R package. Phosphopeptide sequences were matched to protein sequences to determine phosphosite positions. If multiple phosphopeptides mapped to the same phosphosite, we selected the site with the higher mean signal in control samples. Phosphosite readouts for kinases directly targeted by the inhibitors we used were taken from the SIGNOR 2.0 database [40]. Posttranslational modification set enrichment analysis (PTM-SEA) was performed using the ssGSEA 2.0 R script (ssgsea-cli.R) and the PERT-, PATH- and KINASE-signature categories of the PTMsigDB v1.9.0 database [41] and kinase/phosphatase signatures derived from SIGNOR. We used fold-change-signed log10-transformed FDR values from mapDIA as input scores and protein-centric phosphosite positions as identifiers. Peptides with multiple phosphorylated residues were demultiplexed as suggested in the original publication. Normalized enrichment scores (NES) and global false-discovery-rate-adjusted p-values (FDR) were calculated with the number of permutations set to 100000, “area.under.RES” as the test statistic, a weight of one, no additional normalization and a minimum overlap of two measured sites per signature. Heatmaps for visualization of phosphoproteomics results were generated using the Complex-Heatmap 2.9.0 R package [25]. The numbers of shared differential abundant phosphopeptides were visualized in R with the UpSetR package.

### Kinase signaling network analysis

Network analysis and visualization was performed with the igraph 1.3.2 [42], tidygraph 1.2.1 and ggraph 2.0.5 R packages. We mapped normalized enrichment scores of kinase signatures from PTMsigDB to nodes and phosphosite log-fold-changes to edges of the global human SIGNOR 2.0 signaling network (retrieved on 23.04.2021) [40]. Perturbation subnetworks were identified from perturbed nodes (kinase signature FDR <= 0.05) and perturbed edges (phosphosite FDR <= 0.05). Briefly, we calculated all shortest paths with a maximum length of two edges between perturbed nodes, combined them with perturbed edges and filtered for the largest connected component. The subnetworks were further refined by pruning them of redundant shortest paths according to whether they could be annotated with measured phosphosites. We further combined the resulting treatment-specific subnetworks into PDO-specific networks, calculated degree and eigenvector centralities of all nodes and tested for overrepresentation of coding mutations using a two-sided Fisher’s Exact Test.

### Imaging mass cytometry

Four μm tissue sections were placed on silane-coated glass slides, dried overnight at 37 °C and stored at 4 °C. Carrier-free IgG antibodies (Table S2) were conjugated to purified lanthanide metals with the MaxPar antibody labeling kit and protocol (Fluidigm) as described by Ijsselsteijn et al [43]. Imaging mass cytometry immunodetection was performed following the protocol described previously by Ijsselsteijn et al [43]. In short, tissue sections were deparaffinized through immersion in xylene for 3 times 5 minutes and rehydrated in decreasing concentrations of ethanol. 10x low pH antigen retrieval solution (Thermo Scientific, #00-4955-58) was diluted in purified water and preheated for 10 minutes in a microwave. The sections were rinsed in unheated 1x low pH antigen retrieval solution, boiled for 10 minutes in the preheated buffer and cooled down to room temperature for 1 hour. Sections were rinsed with PBS-TB (PBS supplemented with 0,05% Tween and 1% BSA) and incubated for 30 minutes with 200μl Superblock blocking buffer (Thermo Scientific, #37515). Antibody incubation was split into two steps: a 5 hour incubation at room temperature and an overnight at 4 °C incubation. The antibody mix for the 5 hour incubation was prepared by diluting the first half of antibodies (Table S2) in PBS-TB after which 100μl of antibody mix was added to the tissue sections and incubated for 5 hours at room temperature in a humid chamber. The sections were washed three times for 5 minutes with PBS-TB and incubated overnight at 4 °C in a humid chamber with the remaining antibodies (Table S2) diluted in PBS-TB. The sections were washed three times with PBS-TB and incubated for 5 minutes at room temperature with 100μl Intercalator Ir (1,25μM diluted in PBS-TB). Tissue sections were washed two times for 5 minutes with PBS-TB and once with purified water for 5 minutes. Finally, the sections were dried under an air flow and stored at RT until ablation. For each tissue, 8 regions of interest were chosen based on hematoxylin and eosin staining on consecutive sections that were representative for the whole tissue. The Hyperion imaging mass cytometry system (Fluidigm) was calibrated using a 3-element tuning slide (Fluidigm) following the manufacturers settings with an extra threshold of 1500 mean duals detected for 175Lu. In total, 40 ROIs of 1000×1000μm were ablated after which 3 ROIs were excluded due to poor quality.

### Imaging mass cytometry data analysis

Data was exported as .MCD files and for each ROI color TIFF images were created containing the DNA, vimentin and keratin signal using the Fluidigm MCD viewer. These were used to segment the images into nucleus, membrane and background using a random forest classifier in Ilastik [44]. The exported probability maps were used to create cell masks in cell profiler [45]. Simultaneously, the MCD files were transformed to .OME.TIFF files using the Fluidigm MCD-viewer. In Ilastik, pixels were assigned to either ‘signal’ or ‘background’ per marker to train a random forest classifier, which was applied to the entire dataset. Data was exported as binary segmentation masks for each marker where the ‘background’ pixels were set to 0 and the ‘signal’ pixels to 1. These signal masks together with the cell masks and ome.tiff files were loaded into ImaCytE [46] to create FCS files containing per cell the mean pixel intensity for each marker. HSNE clustering on the FCS files of all images was performed in Cytosplore [47] to generate phenotype clusters which were mapped back onto the images in IMACyte. Each cluster was visually confirmed using the original .MCD files and combined when similar clusters were observed. The phenotype clusters were further processed using R to calculate cell densities (cells/mm2), composition of present immune cells and to investigate the expression of selected markers of interests in specific phenotype clusters for each sample. Threshold of 10% was set for the marker values, i.e. a cell is considered as positive for a certain marker if in at least 10% of its area the marker was positively detected. The Spearman pairwise correlation heatmaps of cell phenotypes were also calculated for each sample separately. To assess the spatial organization and cell-cell interactions Voronoi diagrams were created and all cell-cell interactions (direct neighbors) were counted. To test whether the number of direct interactions of each pairwise cell type combination is significantly different than expected by chance, a Monte Carlo simulation with 1000 iterations was performed in which the location of the cells from the imaged slide was randomly permuted, while keeping the number of cells from each type constant and the overall cellular positions fixed. Then a z-score and p-value was calculated to assess avoidance or attraction (z-score < 2, z-score > 2, p < 0.01). A combined cohort z-score was then calculated using the Stouffer’s z-score method for overall meta analysis and plotted as heatmaps (Figure S6). PD1+/PDL1+ microaggregates (min. 1 PDL1+ tumor cell and min. 1 PD1+ immune cell) were identified by finding connected components in a graph crated from the Voronoi diagram. These microaggregates were color marked with the respective cell type colors and plotted as Voronoi diagrams (Figure 6A). To test if the number of PD1+/PDL1+ microaggregates per tumor was significantly different than expected by chance, we again performed a Monte Carlo simulation as described above and calculated z-scores and p-values (Table S3).

## Results

### Proteogenomic characterization of a living biobank

We generated and molecularly characterized a living biobank using organoid technology [48] (Supplementary Figure 1A) as the basis for the subsequent analyses. Tumor samples from 15 CRC patients, including microsatellite instable (MSI) and microsatellite stable (MSS) tumors were used to generate PDOs which were then characterized using exome-sequencing, transcriptome-sequencing, and SWATH-MS-based proteomics. Genomic characterization including CRC driver genes [49] and a panel of immune-related genes frequently mutated in CRC [50] reaffirmed previously reported somatic alterations [50] and showed that the biobank is representative of CRC (Figure 2A, Supplementary Figure 1B). As expected, MSS PDOs have few targetable mutations whereas mutations in immune-related genes including mutations in Human Leukocyte Antigen (HLA) class I and class II genes were almost entirely detectable in MSI PDOs (Figure 2A). WNT signaling (*CTNNB1* (ß-catenin)) and TGFß signaling genes (*SMAD2, SMAD4*) were mutated in both subtypes. We then predicted the presence of potential neoantigens including neoantigens derived from missense mutations and from fusion genes using RNA-seq data and our recently developed tool for predicting neoantigens [18] (Figure 2A). Both, HLA class I and class II-associated neoantigens from gene fusions were mostly detectable in MSS PDOs. Analyses of the neoantigen landscape showed that neoantigens from tandem duplications from acid ceramidase (*ASAH1*) were predicted in 25% of the PDOs, suggesting that this public neoantigen can be used for developing therapeutic vaccination using off-the-shelf vaccine for this CRC cohort.

**Figure 2.**
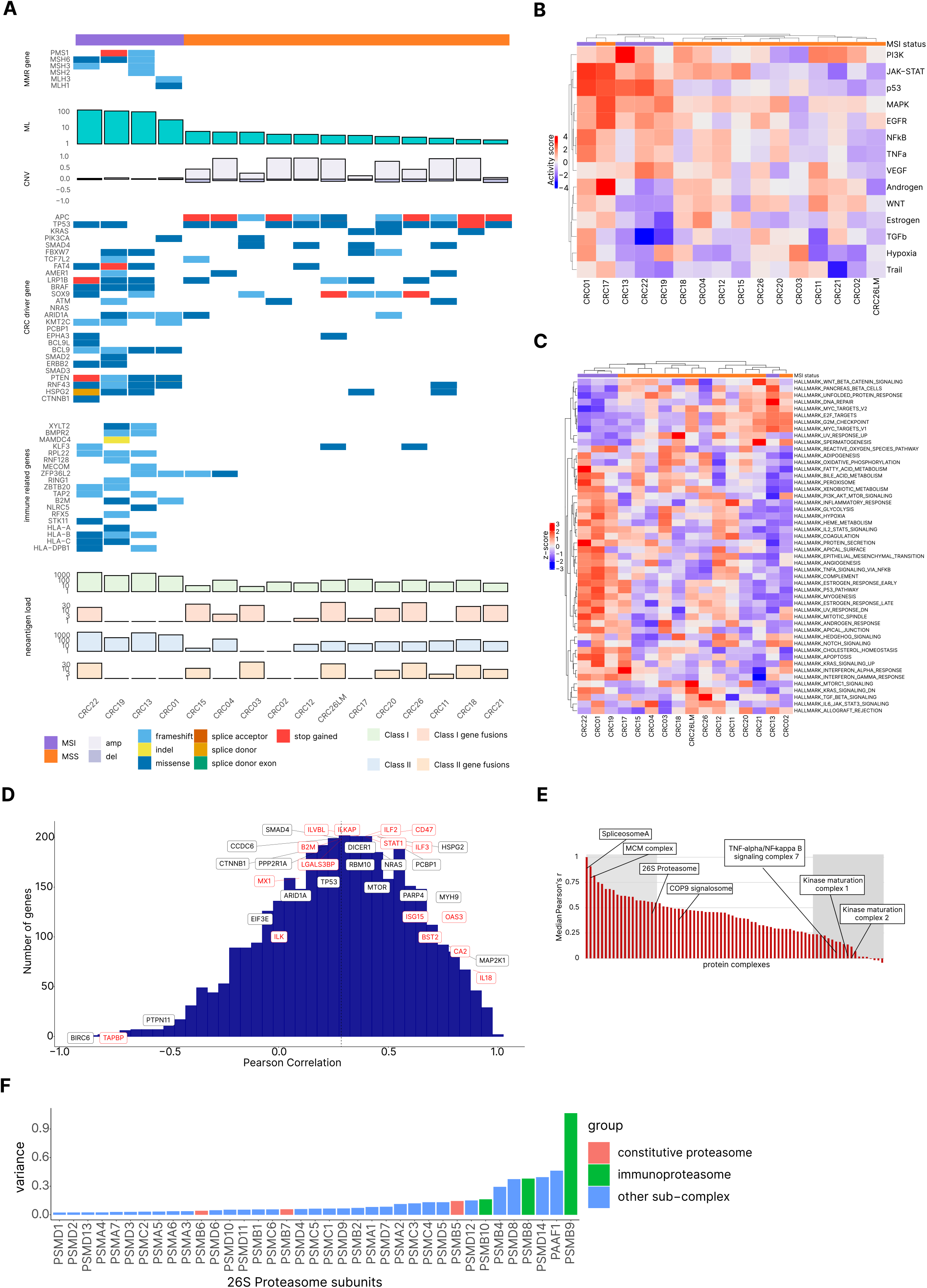
Proteogenomic analysis of PDOs from CRC patients. **(A)** Genetic profiles of the PDOs ordered according to the mutational load (ML). MMR: mismatch-repair. ML: mutational load. CNV: copy number variation. **(B)** Analysis of the cancer pathways of the PDOs using bulk RNA-seq data and PROGENy. The pathway activity scores are z-scaled and clustered hierarchically by euclidean distance and complete linkage. **(C)** Pathway analysis of the hallmark gene sets from MSigDB of the PDOs using proteomic data (SWATH-MS). The heat map shows z-scores of enrichment scores derived from Gene Set Variation Analysis (GSVA) and clustered hierarchically by Pearson correlation as distance metric and complete linkage. **(D)** Correlation analysis between RNA-seq data and proteomics data. The histogram shows gene-wise Pearson correlation between transcriptome and proteome levels. Denoted are driver genes (black) and immune-related genes (red). The average gene-wise Pearson correlation is 0.29 (dashed line). **(E)** Protein complexes ranked according to the co-abundance observed for complex members. Shaded areas, left: stable complexes (top 25%), right: variable complexes (bottom 25%). MCM: mini chromosome maintenance. COP9: constitutive photomorphogenesis 9. **(F)** Variance of the protein levels of the 26S proteasome subunits across all PDOs.

Analysis of the steady-state transcriptomic data revealed expression of genes involved in chemokine-mediated signaling pathways, indicating possible crosstalk with immune cells (Supplementary Figure 1C). We then assembled a panel of genes associated with cancer immunity and immune evasion mechanisms including checkpoint molecules, antigen processing and presentation genes, specific chemokines and cytokines, and tumor cell-specific interferon γ-related (IFN_γ_) genes [51]. The expression of these genes was highly heterogeneous in the MSI and MSS samples with HLA class I genes showing increased expression in MSI relatively to MSS PDOs (Supplementary Figure 1D). Analysis of cancer pathways using transcriptomic data showed heterogeneous pathway activity and partitioning of the profiles in two main subgroups, one of which was predominantly MSI subtype related (Figure 2B).

Analysis of the protein expression data following SWATH-MS of the PDOs showed 3723 unique proteins across the samples (Supplementary Figure 1E). Pathway analysis using protein expression data showed partitioning related to the MSI and MSS subtypes (Figure 2C) with notable exception of CRC13 from the MSI group and CRC17 from the MSS group. Within the MSI group there was a tendency for coordinated upregulation of the immune-related pathways, IFN_α_ response, IFN_γ_ response and IL2/STAT5 (Figure 2C). Gene-wise correlation analysis between RNA and protein expression (average gene-wise Pearson correlation 0.29, Figure 2D) showed that RNA expression is a poor predictor of protein expression for CRC driver genes and for immune-related genes.

As proteins generally exert their function in coordination with other proteins in cellular pathways complexed with other proteins [52,53], we analyzed the co-abundance for complex members of 78 complexes with at least 5 protein members across all PDOs (Table S4). Complexes with variable subunit composition included *TNFα/NFκB* signaling complex 7 and kinase maturation complexes 1 and 2 (Figure 2E). In our analysis, complexes with invariant subunit composition included spliceosome-A, mini chromosome maintenance (MCM) complex and 26S proteasome. However, within the 26S proteasome, large variability was observed for the protein complex subunits PSMB9 and PSMB8 of the immunoproteasome (Figure 2F), the expression of which is associated with immune response to ICBs in melanoma [54]. Noteworthy, the expression of PSMB9 was decreased in a large fraction of the organoids (Supplementary Figure 1F).

In summary, steady-state multi-omics profiling of the PDOs revealed molecular heterogeneity within the clinical subgroups of MSI and MSS tumors and showed a number of altered signaling pathways that could determine cellular responses to drug treatment. However, based solely on these multi-omic data, the identification of the mechanisms that shape key tumor vulnerabilities and determine response to targeted therapy remained elusive.

### Functional precision profiling of PDOs reveals off-target effects and pathway crosstalk

In order to investigate the effects of the targeted drugs on specific pro-tumorigenic pathways as well as to identify potential signaling crosstalk with anti-tumorigenic pathways, we used perturbation experiments and quantitative phosphoproteomic profiling. We carried out perturbation experiments on selected PDO’s using a panel of kinase inhibitors (BRAFi, MEKi, mTORi, PI3Ki, TAKi, TBKi, Supplementary Figure 2A) and one stimulus (TNF_α_) followed by RNA-sequencing and SWATH-MS phosphoproteomics measurements. SWATH-MS phosphoproteomics data of the signaling perturbation experiments was highly reproducible and comprised 10,664 phosphopeptides that were mapped to 7,778 unique phosphosites (Figure 3A, Supplementary Figure 2B). Unsupervised clustering of the identified phosphosites showed that the profiles were mostly specific for PDOs (Supplementary Figure 2C), suggesting patient-specific rather than treatment-specific phosphoproteomes. The overlap of the regulated phosphopeptides between the treatments was in the lower percentile range (Figure 3B) even for treatments with inhibitors targeting kinases in the same pathway (e.g. the inhibition of BRAF and MEK in the MAPK pathway).

**Figure 3.**
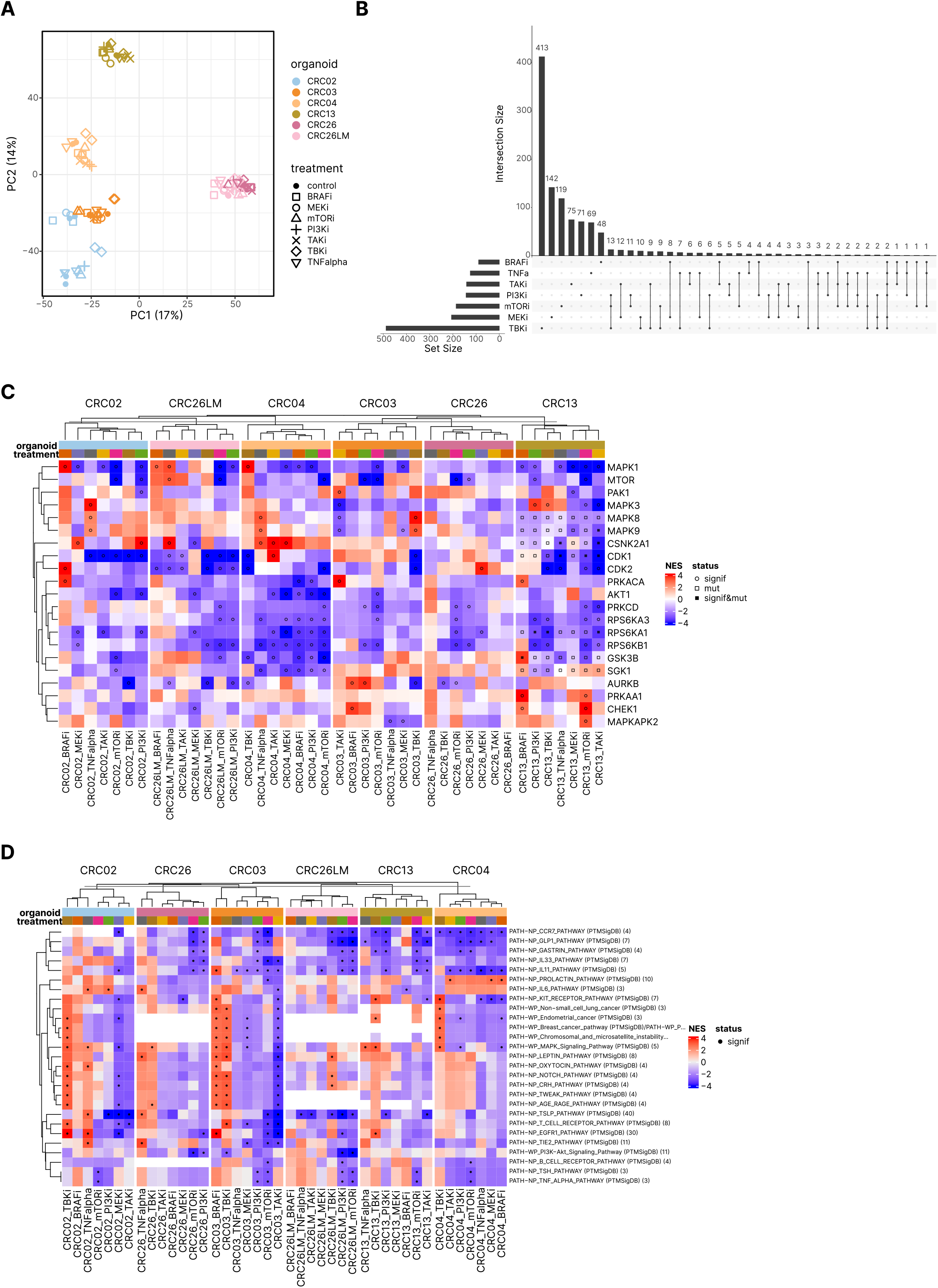
Functional profiling experiments of the PDOs with targeted drugs. **(A)** PCA of the phosphoproteomic data. **(B)** UpSet plot of regulated phosphopeptides (|log2FC|>1, FDR<0.05) following treatment with specific kinase inhibitors. **(C)** Heatmap of normalized enrichment scores (NES) of phosphorylation signatures from PTMSigDB [41] and SIGNOR [40] with at least five phosphorylation sites representing changes in kinase activities following treatment of PDOs with specific kinase inhibitors or TNF_α_ (FDR<0.05), clustered by complete linkage of Euclidean distances. **(D)** Normalized enrichment scores (NES) of phosphorylation signatures representing changes in pathway activities following treatment of PDOs with specific kinase inhibitors or TNF_α_ (FDR<0.05, database and number of phosphorylation sites shown in brackets).

We then analyzed 103 kinases using 508 phosphosites that matched phosphosites in a curated database of phosphosite-specific signatures (PTMsigDB) [41] (Table S5). The analysis revealed highly diverse responses to the perturbations in the PDOs and showed reduced activities of the targeted kinases including MAPK1, AKT1, CDK1, CDK2 upon treatment of the PDO’s with the respective inhibitors (Figure 3C, Supplementary Figure 2D). However, there were extensive off-target effects of the inhibitors including activation of kinases in non-targeted cascades that were specific for both, PDOs and kinase inhibitors. For example, we observed an activation of mTOR following BRAF and TBK inhibition or activation of AURKB following mTOR inhibition in several PDOs (Figure 3C). Set enrichment analyses with the phosphosite-specific signatures (PTMsigDB) [41] revealed pathway crosstalk with a number of pathways including immune-related pathways, like IL-2, IL-6, and IL-33 pathway (Figure 3D). The pathway crosstalk was PDO- and inhibitor-specific and included increased and reduced pathway activity without any obvious pattern.

### Rewiring of kinase signaling networks is mutation-independent

The observed off-target activation of kinases in non-targeted cascades and the resulting pathway crosstalk necessitates detailed characterization of the signal transduction network in order to identify optimal targets for effective modulation of the respective pathways. Signal transduction networks are highly adaptable and dynamic, the properties of which are primarily determined by the network topology [55]. In an attempt to reconstruct signaling networks in individual patients we developed a computational method using the phosphoproteomic data and *a priori* knowledge of protein-protein interaction networks (see Methods). Briefly, we assigned kinase activities to nodes and phosphosites to edges of the SIGNOR 2.0 signaling network [40]. To identify subnetworks probed by the perturbations with kinase inhibitors only nodes with differential kinase activity based on the enrichment score calculated using PTMSEA [41] and edges differentially phosphorylated were considered, and the largest module was extracted (see Methods). The individual subnetworks resulting from each perturbation were then amalgamated into a combined kinase signaling network for the corresponding PDO. The kinase signaling networks showed a large extent of heterogeneity with varying numbers of nodes and edges as well as kinase activities and target site phosphorylations (Figure 4A, Supplementary Figure 3). Strikingly, rewiring of the kinase signaling networks was largely independent of the mutations. We found no significant association of mutations with PDO networks (CRC02 pval=0.18, CRC03 pval=0.63, CRC04 pval=1.00, CRC13 pval=0.61, CRC26 pval=0.26, CRC26LM pval=1.00, Fisher’s exact test;). In the 5 MSS PDOs harboring between 116 and 230 mutations (coding variants), there were zero (CRC03, CRC26LM), one (CRC04, CRC26) and two nodes (CRC02) with mutated proteins. Even in the MSI PDO (CRC13) with a large number of mutations (2850 coding variants) there was no significant association of the mutations with the edges in the signaling kinase network (pval=0.61).

**Figure 4.**
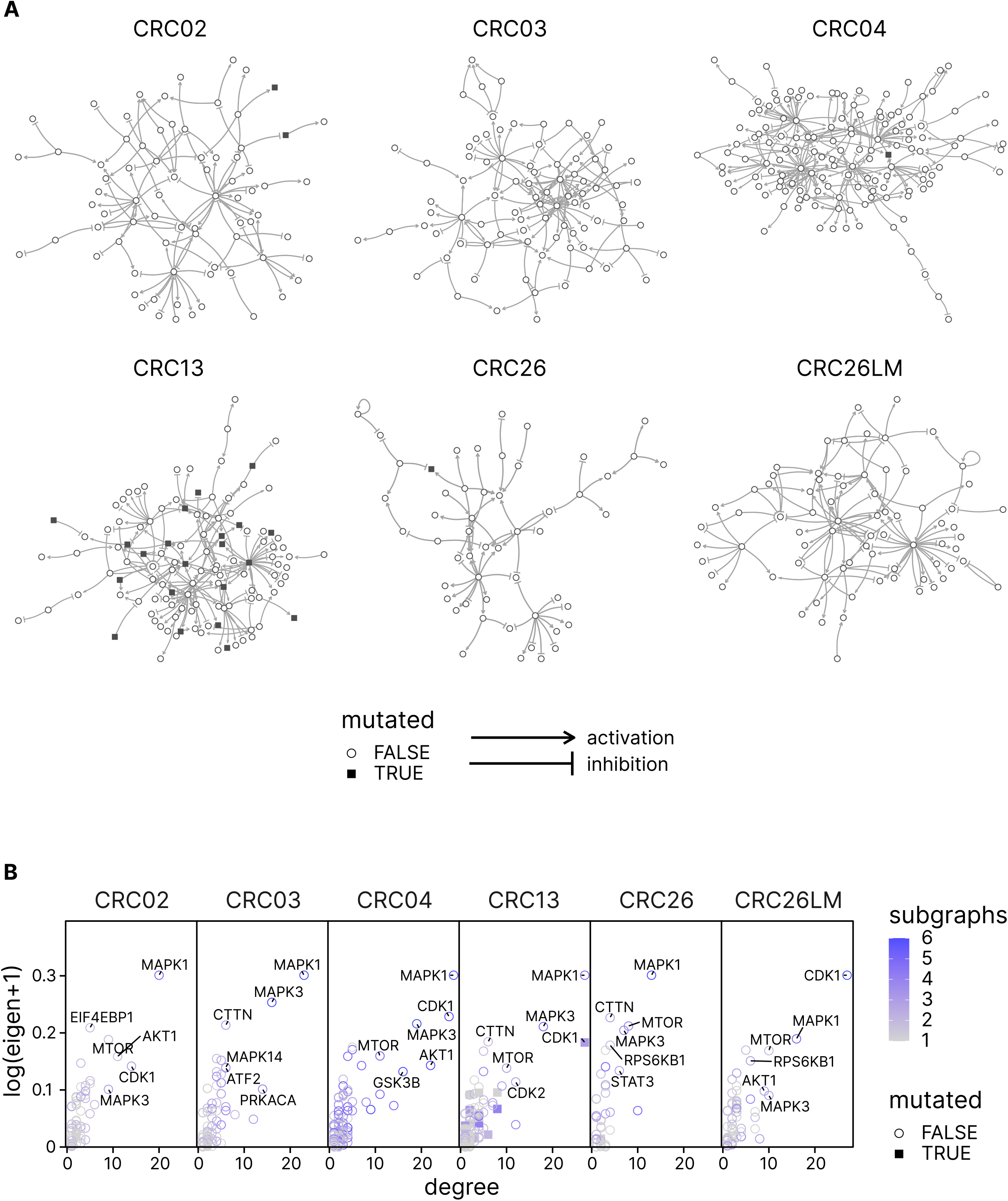
Comparative analysis of the kinase network topologies for the perturbed PDOs. **(A)** Visual representation of the reconstructed kinase networks. Highlighted in blue are kinases harboring mutations. **(B)** Betweenness and degree centrality measures of kinase nodes in the networks shown in **(A)**.

It has been previously shown that node centrality values are correlated with the importance in maintaining network integrity [56]. We therefore performed comparative analysis of the kinase signaling networks for the PDOs and obtained centrality values for each kinase node. (Figure 4B). As expected, high centrality values were observed for targeted kinases (e.g. mTOR) and for the kinases in the central EGFR-RAS-MAPK pathway (e.g. MAPK1). However, additional nodes like PRKACA or PTPN7 had high centrality values, further supporting the notion of extensive off-target effects and pathway crosstalk.

Overall, using quantitative phosphoproteomic data from PDOs perturbed with kinase inhibitors we were able to reconstruct kinase signaling networks at the patient level. Albeit heterogenous between PDOs, the kinase signaling network topologies revealed intrinsic rewiring that was independent of the harboring mutations.

### Single-cell analysis shows phenotypic and differentiation heterogeneity of PDOs

Our finding of non-mutational rewiring of kinase signaling networks suggests that non-genetic mechanisms like phenotypic plasticity and differentiation status of the tumor cells might have a larger impact on adapting the signaling circuitry and leading to disease phenotypes. According to the cancer stem cell hypothesis, tumors are organized in cell hierarchies similar to normal tissues, with stem cells at the apex, giving rise to transient amplifying (TA) progenitor cells that undergo differentiation into several cell lineages [57]. The revised cancer stem cells model postulates that cancer cells can dynamically shift between a differentiated state and a stem-like state [58], which in CRC is tightly linked to changes in Wnt-signaling. Noteworthy, disrupted differentiation is integral to colon carcinogenesis and is a regulator of cellular plasticity. Most CRCs are diagnosed as moderately differentiated [59] with some cell types being implicated in therapy response. For example, it has been shown that enteroendocrine progenitors support *BRAF*-mutant CRC [60]. Hence, it appears that the hierarchically organized tumor cell heterogeneity and cell plasticity play key roles in both, CRC progression and therapy response as shown recently [61]. We therefore aimed to determine the hierarchically organized tumor cell heterogeneity in our PDOs and employed single-cell RNA sequencing.

We generated transcriptomic profiles from 37,924 cells (after quality control and filtering) which clustered according to the PDOs with CRC26 (primary tumor) and CRC26LM (liver metastasis) being closest (Supplementary Figure 4A). We identified 8 clusters (Figure 5A) which were annotated using curated panels of genes (Figure 5B, Supplementary Figure 2B-C). All PDOs included moderately differentiated stem-like cells, transient-amplifying-(TA)-like cells, goblet-like cells and M-like cells according to the respective markers (Figure 5C, Supplementary Figure 2B-C). The PDOs were highly heterogeneous with respect to the fractions of different cell types, with TA-like cells being most and M-like cells being least abundant (Figure 5C). Analyses of RNA velocity indicated cell hierarchies according to the cancer stem cell hypothesis, with stem cells at the apex, giving rise to TA progenitor cells that undergo differentiation into several cell lineages (Figure 5A). Using a compendium of pathway-responsive gene sets we then assigned activities of 14 canonical cancer pathways to the individual cell types. The cancer pathway activities were variable between the secretory cells and other cell types, and between the PDOs (Figure 5D).

**Figure 5.**
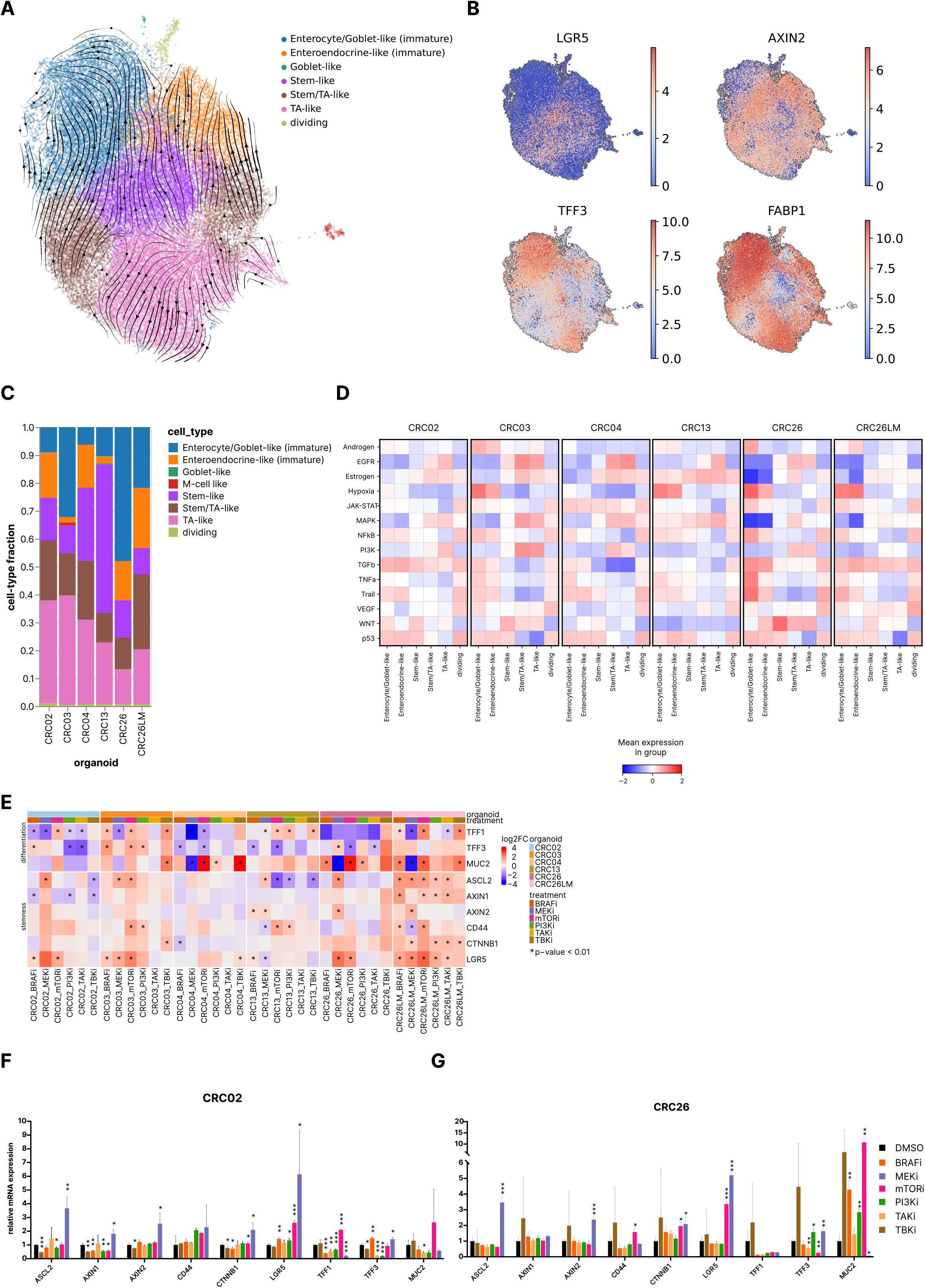
Single cell analysis of PDOs from CRC patients. **(A)** UMAP plot of batch-corrected scRNA-seq dataset from all organoids, colored by cell-type. RNA velocity vectors are projected on top of the UMAP plot. **(B)** UMAP plot from (A) colored by gene expression (log(CPM)) of the markers for stem cells (*LGR5*), Wnt target (*AXIN2*), goblet cells (TFF3), and enterocytes (*FABP1*). **(C)** Cellular composition of the PDOs as measured by scRNA-seq. **(D)** Analysis of cancer pathways activation in specific epithelial cell types using PROGENy [23]. **(E)** Heat map of regulation of stem cell markers and differentiation markers following treatments with different kinase inhibitors for 72 hours and qPCR measurements. Each drug treatment was performed in triplicates. Fold change in expression of target genes was calculated by 2−ΔΔCT method [72] using DMSO control for normalization, and GAPDH as an endogenous control. * p < 0.01; unpaired two-tailed Student’s t-test. **(F-G)** Relative mRNA expression of CRC02 and CRC26 organoids from the analysis shown in (E). Error bars indicate + SD and p-values show significant differences between the treated vs DMSO control.* p < 0.05, ** p < 0.01, *** p < 0.001, unpaired two-tailed Student’s t-test.

**Fig. 6.**
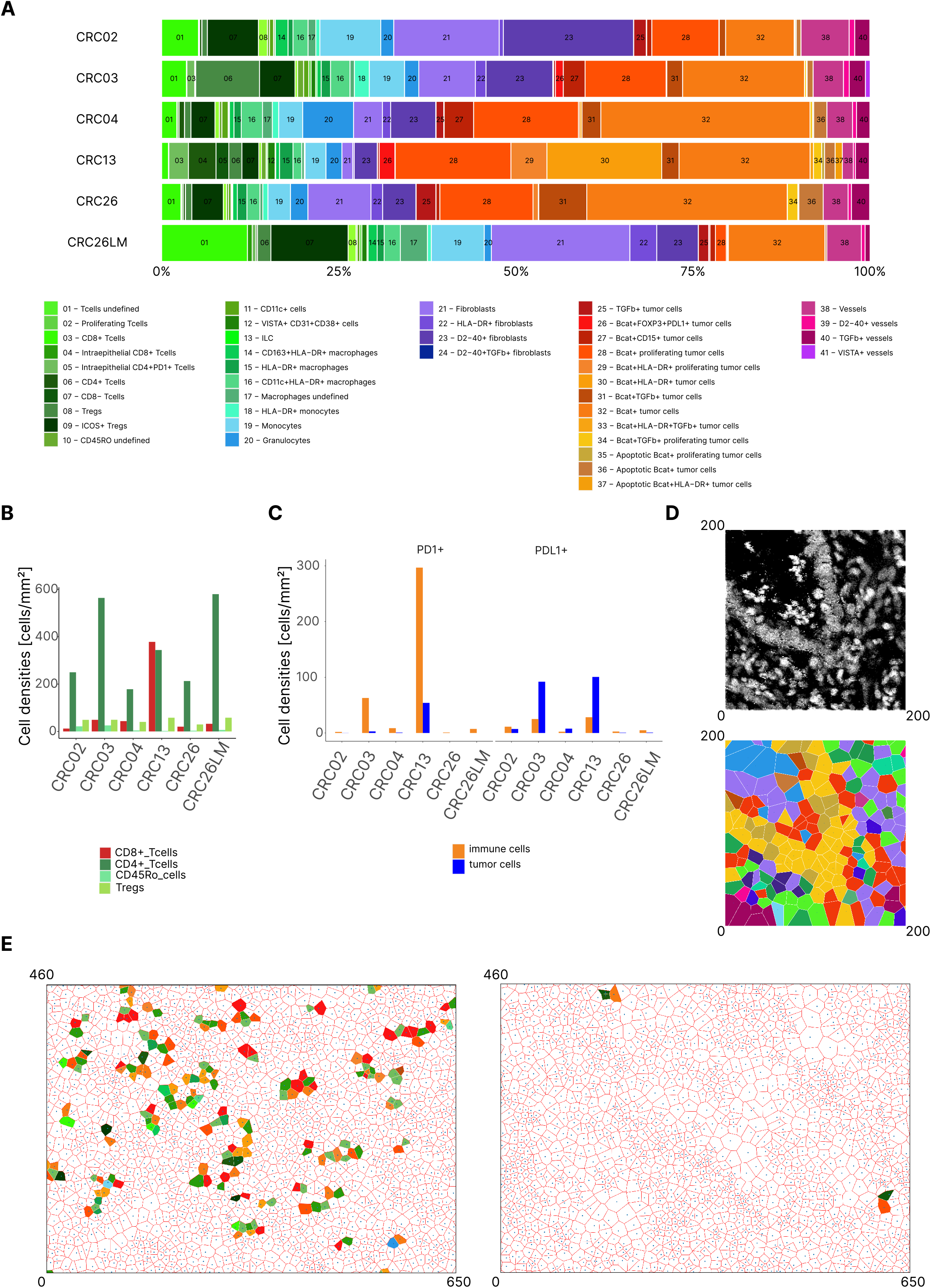
Spatial proteomics of tumor samples using imaging mass cytometry. **(A)** Cellular composition of the TME using 41 cell phenotypes from six tumor tissues from the respective patients. **(B)** Cell densities of CD8^+^, CD4^+^, CD45RO^+^ and Tregs. **(C)** Cell densities of PD1^+^ tumor cells and PD-L1^+^ immune cells. **(D)** Example subsection (200×200 μm) of cell neighborhood analysis using Voronoi diagrams. Upper panel: original image. Lower panel: map following cell phenotype identification and building of Voronoi diagrams. **(E)** Example subsection (200×200 μm) of the cell neighborhood analysis for PD1^+^ tumor cells and PD-L1^+^ immune cells interactions in CRC 13 (left) and CRC03 (right).

### Kinase inhibitors modulate stemness and differentiation pathways

The phenotypic and differentiation plasticity of the tumor cells shown here can have profound effects on the tumor formation, malignant progression, and response to therapy. For example, it has been shown that MEK inhibitors activate *WNT* signaling and induce stem cell plasticity in CRC [61]. Similarly, an experimental evidence was provided showing that therapies targeting the MAPK pathway can redirect developmental trajectories of CRC and can be associated with therapy resistance [62]. However, both studies focused on the inhibitors targeting kinases within the canonical MAPK pathway, i.e. MEK and EGFR/BRAF/MEK. Given the extensive pathway crosstalk in our PDOs, we asked what is the effect of other inhibitors on the phenotypic and differentiation plasticity. We therefore carried out experiments and treated the PDOs for 72 hours with the used kinase inhibitors (BRAFi, MEKi, mTORi, PI3Ki, TAKi, TBKi), and analyzed markers for stemness and differentiation. The results show heterogeneous effects of the MEKi and BRAFi confirming previous studies [61,62]. Moreover, these effects were also observable for other inhibitors including mTORi, PI3Ki, TBKi, and TAKi (Figure 5E). For example in CRC02 MEKi induced upregulation of the stemness marker *LGR5* (Figure 5F), whereas in CRC26 mTORi induced both, upregulation of *LGR5* and the differentiation marker *MUC2* (Figure 5G). Similar diversity was observed for other kinase inhibitors (Figure 5F-G) and other PDOs (Supplementary Figure 2E-H).

### Quantifying heterocellular signaling crosstalk using spatial single-cell proteomics profiling of tumors

Several PDOs showed upregulation of immune pathways following treatment with specific kinase inhibitors, suggesting possible synergistic effects of kinase inhibitors with immune checkpoint blockers. An effective antitumor response following combination therapy of kinase inhibitors and anti-PD-1 or anti-PD-L1 antibodies requires both the presence of CD8+ T cells in the TME as well as CD8+T cell-tumor cell interactions. We therefore used IMC-based multidimensional imaging of the tumor samples to quantify the densities of immune cell subpopulations and identify heterocellular interactions. We previously developed and evaluated a panel of antibodies for IMC (Table S2) on formalin-fixed, paraffin embedded (FFPE) samples for a comprehensive overview of the TME and cancerimmune cell interactions [43], including lineage and functional immune cell markers, surrogates of cancer cell states (proliferation, apoptosis) and structural markers (epithelium, stroma, vessels) (Supplementary Figure 5A). We used this panel and FFPE samples from five primary tumors and one liver metastasis of the CRC patients. Following data-driven identification of single-cell phenotypes (Supplementary Figure 5B), segmentation, and image analysis, we identified 197,454 cells and quantified the densities of five major classes: myeloid, lymphoid, epithelial, fibroblasts, and endothelial cells, which could be further granulated into 41 different cell types (Figure 6A). The cell densities were highly heterogeneous, with CD8^+^ T cells being the most abundant in MSI CRC (Figure 6B, Supplementary Figure 6A). The analyses of the co-expression of immunomodulatory molecules *PD-1, PD-L1, ICOS, LAG3, TIM3*, and *IDO* showed heterogeneous populations of immune and tumor cells (Supplementary Figure 6B). The densities of *PD-1*+ immune cells and *PD-L1*+ tumor cells were highest in tumors from patients CRC03 and CRC13 (Figure 6C), suggesting that these patients are candidates for immunotherapy with anti-PD1/antiPD-L1 antibodies.

In order to generate higher-order information beyond cell densities, we investigated spatial cell-cell interactions. We applied cell neighborhood analysis by defining cell nuclei and associating polygons (Voronoi diagram) to each nucleus (Figure 6D), thereby allowing cells of different sizes and distances to be assessed as neighbors [63]. We used a permutation approach to identify pairwise interactions between cell phenotypes that occurred more or less frequently than expected by chance. This spatial information revealed a number of significant cell-cell interactions, with cell pairs being close neighbors (cell-cell attraction) (Supplementary Figure 6C) or distant neighbors (cell-cell avoidance) (Supplementary Figure 6D). Cellular attractions across all tumors were detectable within the lineages of myeloid, lymphoid, epithelial, fibroblasts, and endothelial cells as well as within the classes (Supplementary Figure 6C). Importantly, neighborhood analysis of *PD-L1*+cells and *PD1*+ cells (defined as direct neighborhood of at least one *PD-L1*+ tumor cell with at least one *PD1*+ immune cell), showed that in patient CRC13 there were *PD-L1*+ tumor cells / *PD-1*+ immune cells interactions whereas in patient CRC03, despite the relatively high densities of both *PD-L1*+ tumor cells and *PD-1*+ immune cells, there were no significant cell-cell interactions (Figure 6C-E, Table S3). Hence, based on the spatial interaction analysis only patient CRC13 would be amenable for a therapy with anti-PD1 or anti-PD-L1 antibodies.

## Discussion

We developed a functional precision oncology approach using PDOs and quantitative phosphoproteomic profiling, and applied this method to demonstrate the feasibility of dissecting tumor cell signaling in individual CRC patients. The information content that can be extracted from these datasets is superior compared to the information content obtained using alternative approaches. Static (i.e. unperturbed) approaches using biopsies or surgical specimens coupled with phosphoproteomic analysis of tumor tissues [64,65] resemble the assessment of the steady-state of the phosphoproteome and are of limited value for inferring kinase signaling networks. Previous functional approaches using phospho-proteomic measurements to construct cancer signaling networks employed cell lines [15,66] and were based on mathematical models that are inherently limited to a small number of molecular interactions [15]. Recently developed platform using *ex vivo* tumor fragments [67] could be a viable alternative to the PDOs, however, given the limited amount of material that can be obtained, the phoshpoproteome coverage is substantially reduced and the number of possible drugs that can be tested highly restricted. Hence, the “next-generation” functional tests shown here enable comprehensive investigation of the intrinsic CRC biology for successfully personalizing treatment.

The results of our functional precision profiling provide new biological insights and have important translational relevance. First, and most importantly, we show that the patient-specific rewiring of the kinase signaling network is unaffected by mutations in CRC. Our results suggest that the responses to targeted therapy are determined by non-genetic mechanisms such as those conveyed by phenotypic plasticity [68]. Single-cell RNA sequencing of the PDOs showed heterogeneous pathway activation in epithelial cell subsets, further supporting the notion of non-genetic mechanisms determining cellular response to drug treatment. We also provide experimental evidence that kinase inhibitors targeting canonical and non-canonical pathways modulate stemness and differentiation pathways, implicating that also re-purposed drugs are re-routing developmental trajectories of CRC. Our finding of mutation-independent signaling rewiring is supported by a growing body of literature suggesting that cancer phenotypes and the responses to therapy are determined by non-genetic mechanisms, in addition to the mutation-driven mechanisms commonly considered. For example, a CRC classification system previously proposed associates epithelial cellular phenotypes like stem-like, Goblet-like or enterocyte cells with responses to cetuximab and standard-of-care chemotherapy [69]. Similarly, recent work using PDX models showed that EGFR inhibition in CRC tumors induces Paneth-like phenotypic rewiring [70], suggesting that cellular plasticity is shaping drug response in cancer. Hence, *in vivo* data using preclinical models and clinical data from large cohorts provide additional evidence for the importance of CRC tumor cell plasticity for the response to targeted therapy. In fact, phenotypic plasticity and disrupted differentiation have been recently proposed as discrete hallmark capability of cancer [68].

Second, we show that kinase inhibitors can induce profound off-target effects resulting in the modulation of both, oncogenic and immune-related pathways. These off-target effects might explain lack of efficacy of targeted therapies as well as failure of combination therapies with immune checkpoint blockers. Off-target effects due to signaling crosstalk, feedback and feedforward mechanisms, as well as signaling network adaptations, have been previously reported in a variety of cancers and model systems [55]. However, predicting such off-target effects of specific kinase inhibitors for individual patients based on static multi-omic measurements is not possible. Hence, information-reach assays based on perturbation experiments and phosphoproteomic measurements as presented here are required.

Third, complementing our functional precision profiling with extrinsic information from histology using IMC of the primary tumors enabled us to quantify spatial heterocellular crosstalk and tumorimmune cell interactions, and hence, provide rationale for combination therapy with immune check-point blockers. Thus, our *tour de force* work based on functional precision profiling and single-cell spatial analysis might serve as a blueprint for developing next-generation functional precision oncology platforms for predicting combination therapy response in individuals with metastatic CRC and possibly also other cancers.

Our work has several limitations that can be addressed in future studies. One limitation of our method for inferring kinase network topologies is the use of literature-mined networks. Literature-mined networks are biased towards well-known kinases and pathways as evident by the number of kinases used in our downstream analyses. Given the limited annotations in the public repositories we were able to use about 10% of the phosphoproteomic data (600 phosphopeptides out of 6000 measured). A promising method for revealing new information on kinase-kinase relationships based on chemical phospho-proteomics was recently published [56] and can be used to infer additional kinase-kinase interactions. Another limitation of our study is the small number of patients and PDOs. Large-scale efforts are needed to investigate big cohorts of patients and potentially identify patterns for patient stratification. Global analysis of the phosphoproteomic data here showed clustering according to patients rather than treatments. Hence, it is intriguing to speculate that there is a limited number of signaling states that could be consequently exploited to stratify patients and ultimately inform therapy. Finally, we used only bulk phosphoproteomic data and could not assign signaling pathways/states to specific epithelial cell subsets. However, technological developments using ultra-high sensitive mass spectrometers are improving [71] and could in near future enable single-cell proteome measurements to gain insights into the cellular heterogeneity.

In summary, the conceptual advances and the insights from the deep molecular and cellular phenotyping we show here challenge the notion that the information flow following kinase inhibition occurs only within specific signaling cascades. We also provide a unique resource of high quality multi-omics and multi-modal data as well as the corresponding living biobank that can be exploited for both, investigation of intrinsic biology of CRC cells as well as the development of novel methods for interrogating intra- and inter-cellular crosstalk. Finally, our multi-modal profiling approach could provide the basis for the development of a platform for informing precision (immuno)oncology in CRC.

## Data and code availability

The processed data supporting the findings of this study (including exome sequencing, RNA-sequencing, proteomics, phosphoproteomics, single-cell RNA-sequencing, high-dimensional TIFF images, single-cell spatial information and phenotypes) are available online at Zenodo (https://doi.org/10.5281/zenodo.7015015).

The MS data which were used to generate the SWATH spectral library, the SWATH raw files and the quantitative results from the SWATH-MS analysis reported in this paper have been deposited in the PRIDE proteomics data repository (https://www.ebi.ac.uk/pride/archive/) under the following accession numbers:

1. Baseline expression experiment: PXD019124;
2. Total cell lysate in cancer vs metastasis: PXD018922;
3. Perturbation expression experiment: PXD018913;
4. Spectral library for phosphopeptides: PXD018862;
5. Spectral library for total cell lysate: PXD018835:

The code used to produce the results of this study is available at https://github.com/icbi-lab/plattner_2022

## Acknowledgments

This work was supported by the European Research Council (grant agreement No 786295 to ZT), by the Horizon 2020 IMI project imSAVAR (grant agreement no 853988 to ZT), by the Austrian Science Fund (FWF) (projects I3291 and I3978 to ZT, and project T974-B30 to FF), by the Vienna Science and Technology Fund (Project LS16-025 to ZT), Österreichische Krebshilfe Tirol (project FF18015 to GL). CP and GS were supported by a DOC-fellowship from the Austrian Academy of Sciences. ZT is a member of the German Research Foundation (DFG) project TRR 241(INF). Antibodies for the Western blots were kindly provided by Jakob Troppmair. We are thankful to Filipp Sokolovski for collecting clinical data.

## Author contributions

ZT conceived the project. GL, RL, AN, SD, NN, NB performed the PDO experiments. CP and AK analyzed the proteomics and phosphoproteomics data. PB and AS carried out proteomics and phosphoproteomics experiments. SS and DÖ recruited patients and performed surgeries. HC, HF, and GH advised the establishment of the living biobank. AN and DW made the PDO stainings. MI performed imaging CyTOF and data analysis. DR and ZL analyzed the imaging CyTOF data. GS analyzed the transriptomics data. DR, GF, and FF analyzed the exome sequencing data and predicted the neoantigens. Anne K prepared sequencing libraries. HF, FG, LAH, RA, NFCCM, and ZT supervised the work. ZT wrote the manuscript with the input from all authors. GL, CP, PB, and AK contributed equally to this work.

## Competing interests

Authors declare no competing interests.

## Supplementary Figure Legends

**Supplementary Figure 1. Genomic and transcriptomic characterisation of PDOs**. (**A)** IF staining of tumor PDOs with Lysozyme (Paneth cells), Chromogranin A (Chr-A) (Enteroendocrine cells), Ki-67 (proliferating cells), MUC2 (Mucin 2) (Goblet cells), EpCAM (arrows point to EpCAM expression, epithelial cells), Phalloidin (F-actin) (brush border), SOX9 and Lgr5+ for (stem cells), and Hoechst (DNA). (Scale bars: 50 μm.). **(B)** Copy number variation analysis for PDOs. Red: amplifications. Blue: deletions. **(C)** Heat map of the steady-state RNA-sequencing data. Eight major clusters were defined and annotated using GO enrichment analysis. The genes are z-scaled and clustered hierarchically by using Pearson correlation as distance and complete linkage. (**D**) Heat map of immune-related genes grouped as antigen presenting cell receptors (APCr), chemokines, chemokine receptors, cytokines, human leukocyte antigens (HLA) and interferon gamma (IFNg). ISG.RS: interferon metagene for IFNG signature genes [73]. (**E**) Heat map of the steady-state proteomics data. Eight major clusters were defined and presented the same way as the gene expressions in (C). (**F**) Heat map representing z-scores of the log2-transformed protein levels of the subunits of the immunoproteasome.

**Supplementary Figure 2. Analysis of the phosphoproteomic data following signaling perturbations**. (**A**) Western blots for selected targets of kinase inhibitors. (**B**) Log-2-fold-changes in the phosphorylation of residues selected as potential downstream targets of kinase inhibitors/activators following the respective treatments (FDR<0.05). (**C**) Heatmap of all log-2-fold-changes in the phosphorylation of residues with FDR<0.05 in at least one treatment, clustered by complete linkage of Euclidean distances.. (**D**) Volcano plots of normalized enrichment scores (NES) for kinase activity signatures from PTMSigDB [41] and SIGNOR [40] following treatment of PDOs with specific kinase inhibitors or TNF_α_.

**Supplementary Figure 3. Signaling networks**. (**A**) Kinase signaling networks for specific PDOs constructed using the phosphoproteomic perturbation data (related to Figure 4A). Log-2-fold changes in kinase activities of significantly perturbed nodes are shown for each treatment.

**Supplementary Figure 4. Single-cell RNA-seq analysis of the PDOs**. (**A**) UMAP of the scRNA-seq data, colored by PDO. (**B**) Violin plots for selected markers. (**C**) Dot plots for the markers used for cell annotation. (D) UMAP of the RNA-sequencing data, colored by cell type. (E-H) Regulation of stem cell markers and differentiation markers following treatments with different kinase inhibitors and qPCR measurements for CRC03, CRC04, CRC13 and CRC26. Each drug treatment was performed in triplicates. Error bars indicate + SD and p-values show significant differences between the treated vs DMSO control.* p < 0.05, ** p < 0.01, *** p < 0.001, unpaired two-tailed Student’s t-test.

**Supplementary Figure 5. Preprocessing of the imaging mass cytometry data**. (**A**) Example IMC images of the major cell types (pseudo coloured) for selected organoids, showing their heterogeneity. Subgroups of major cell types are coloured in similar colour shades. (**B**) Hierarchical clustering of the measured expression values of 34 selected markers used for the phenotyping and recognition of different cell types.

**Supplementary Figure 6. Imaging mass cytometry analyses of the tumors**. (**A**) Cell densities for the four major cell types. (**B**) Cell densities for selected immuno-modulatory molecules. (**C**) Interaction analysis showing cell-cell attraction. Squares are showing the number of PDOs in which significant interactions occur. (**D**) Interaction analysis showing cell-cell avoidance. Squares are showing the number of PDOs in which significant interactions occur.

## Notes

https://doi.org/10.5281/zenodo.7015015

https://github.com/icbi-lab/plattner_2022

### Competing Interest Statement

The authors have declared no competing interest.

### Summary of Updates

Title adaptation and restructuring.

