## Supplementary Figures for "Functional and spatial proteomics profiling reveals intra- and intercellular signaling crosstalk in colorectal cancer"

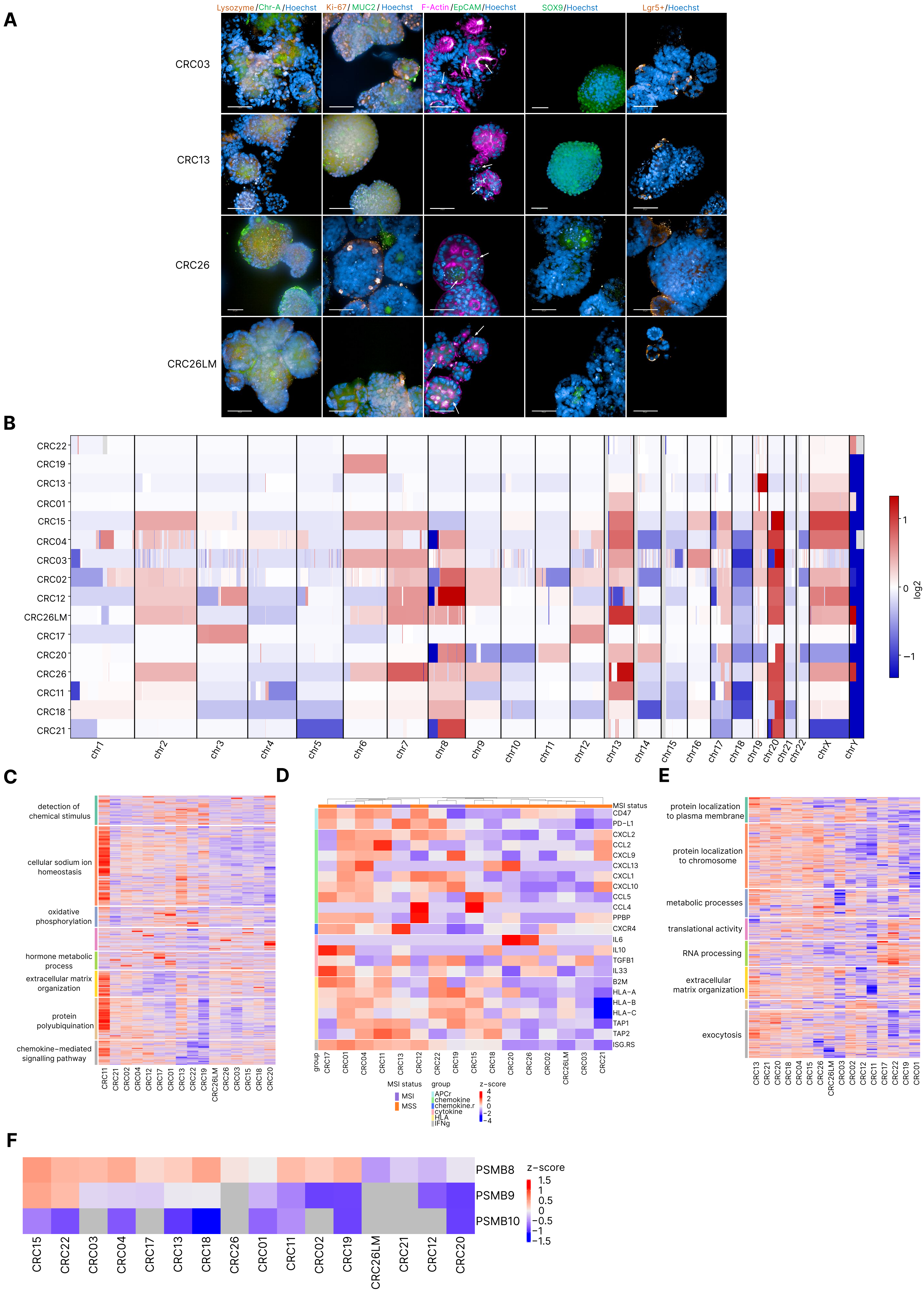


Supplementary Figure 1

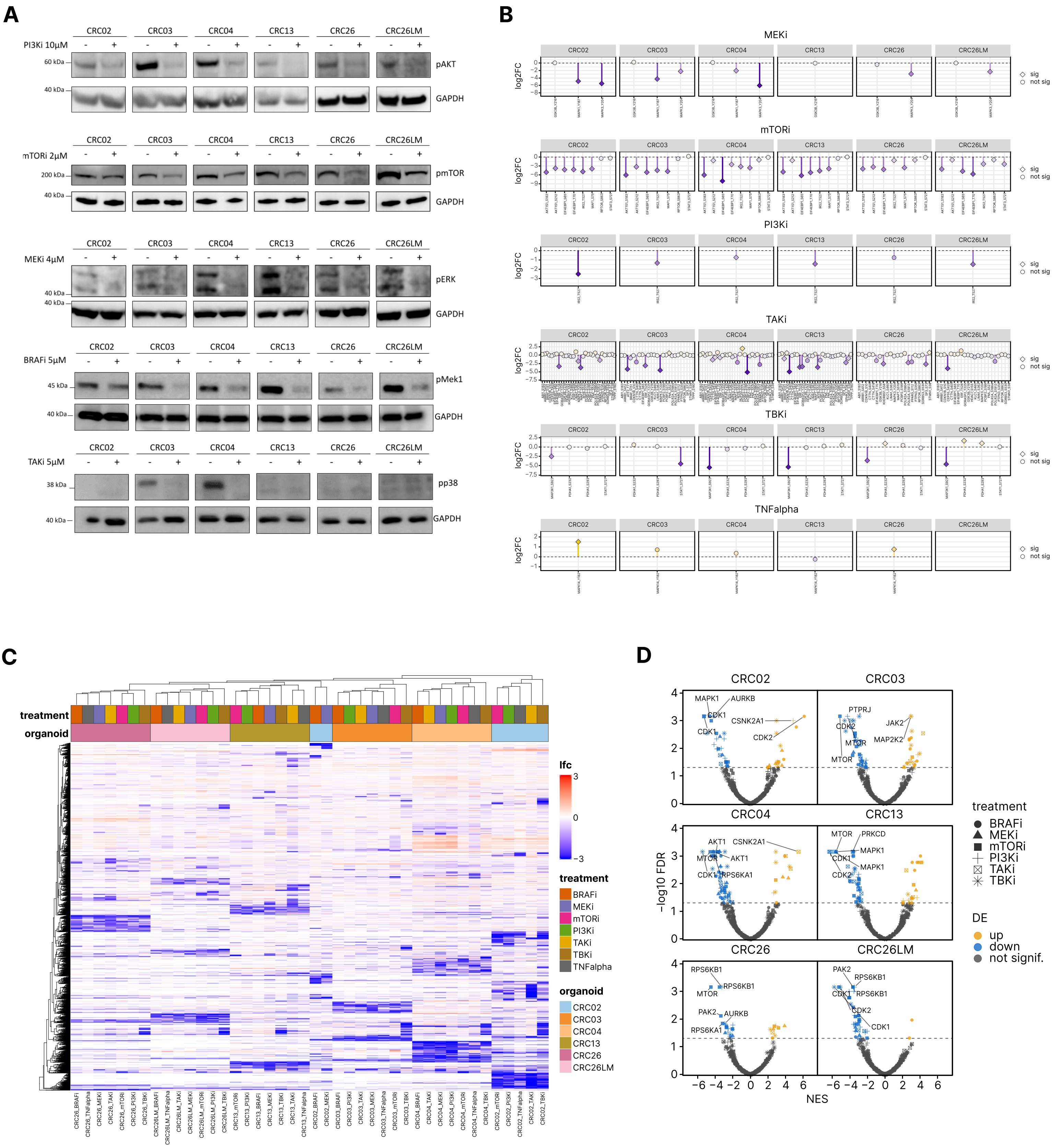


Supplementary Figure 2

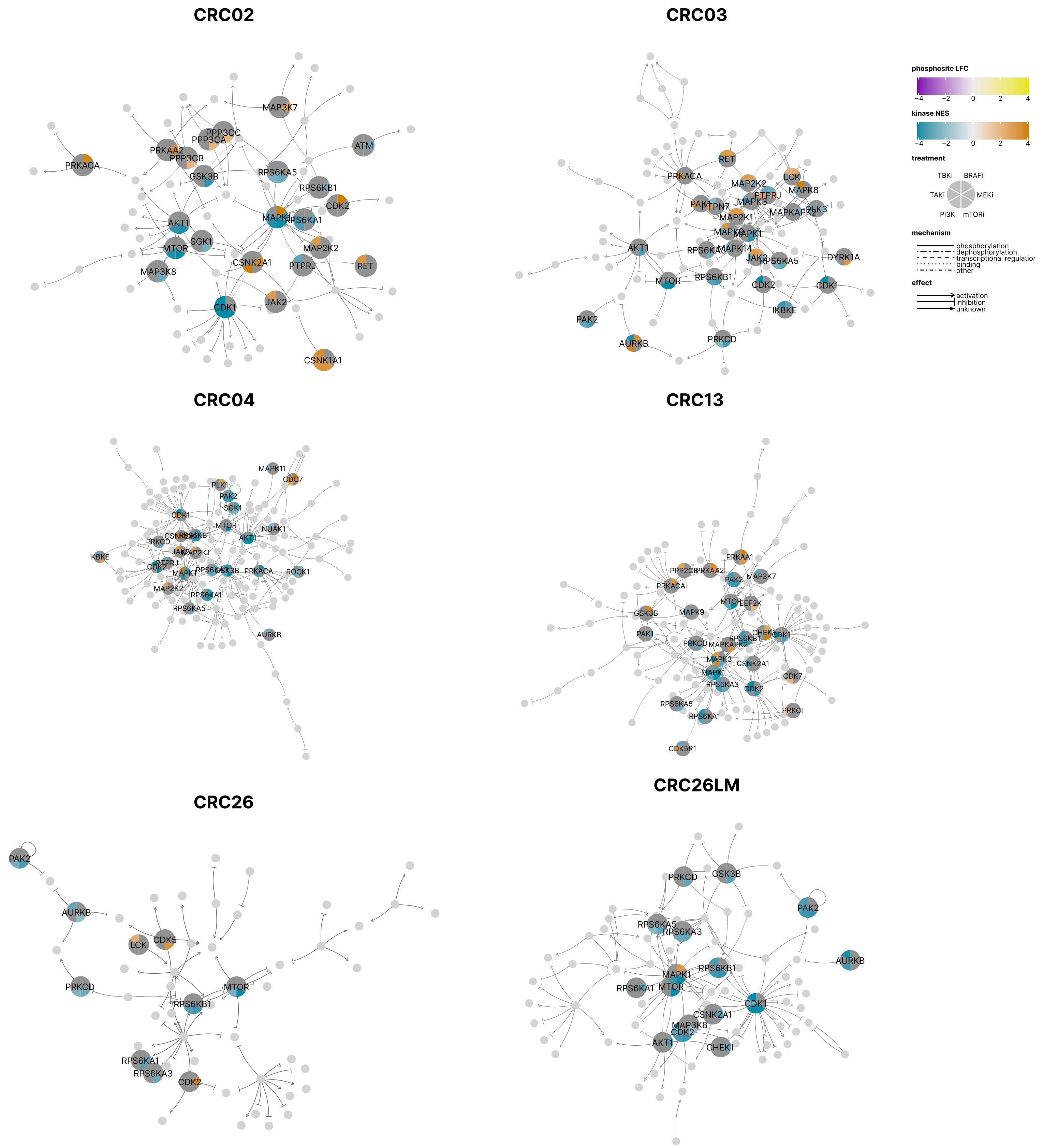


Supplementary Figure 3

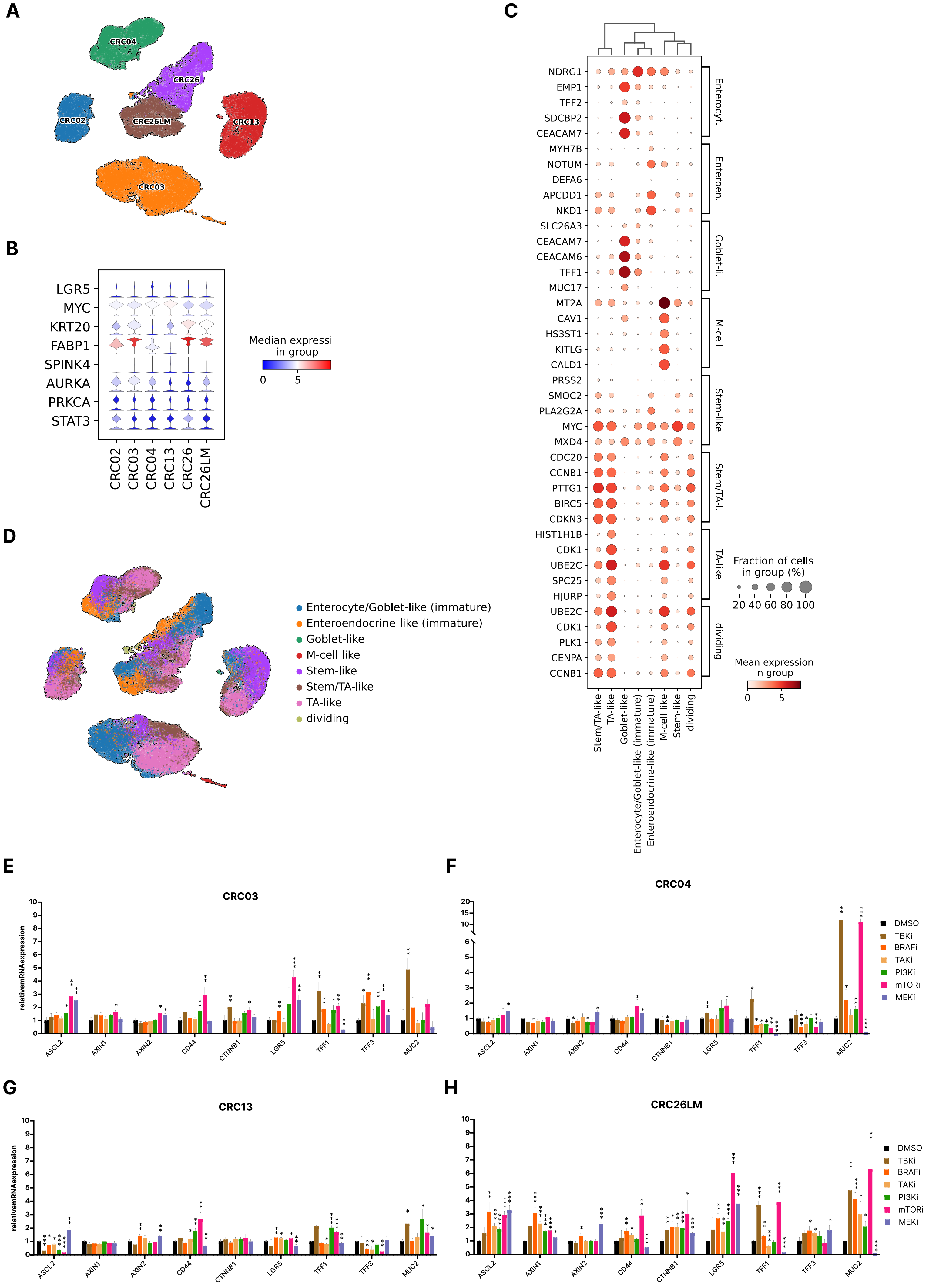


Supplementary Figure 4

**A**

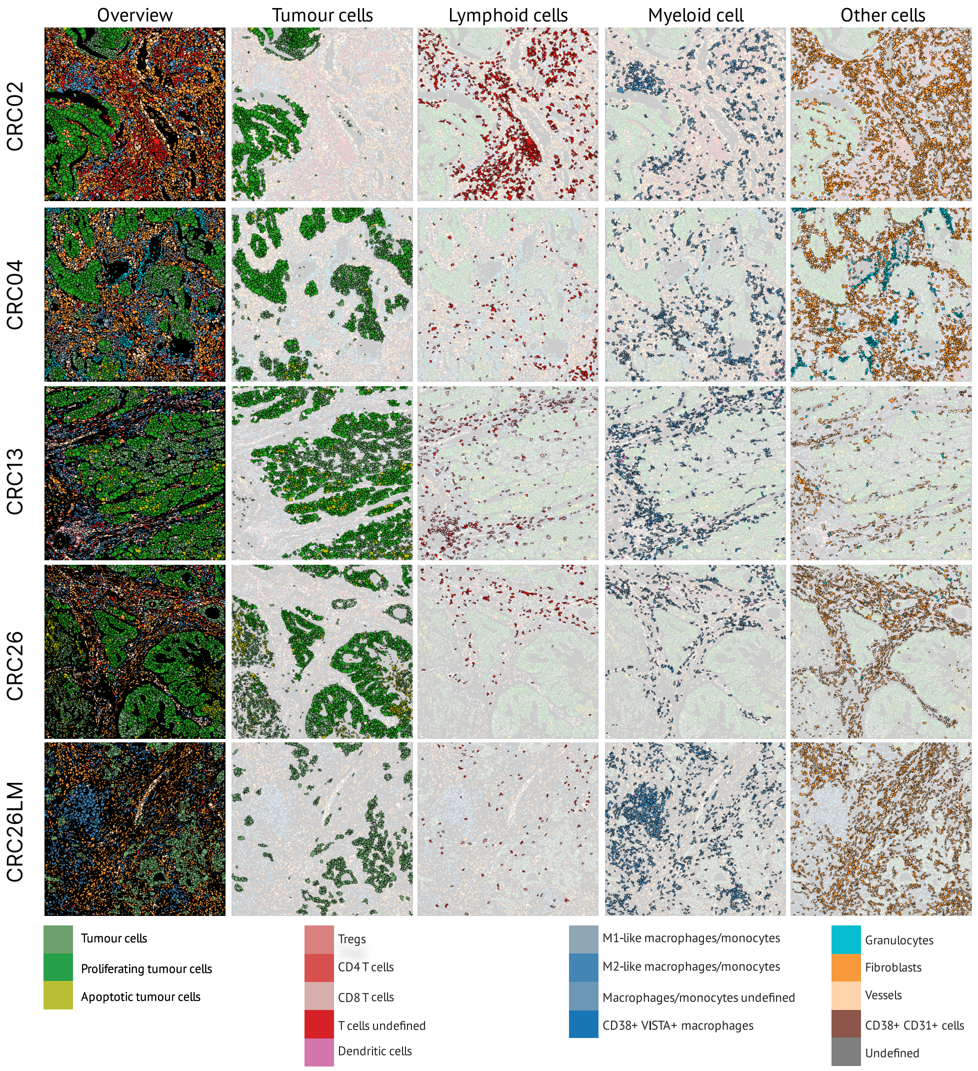


**B**

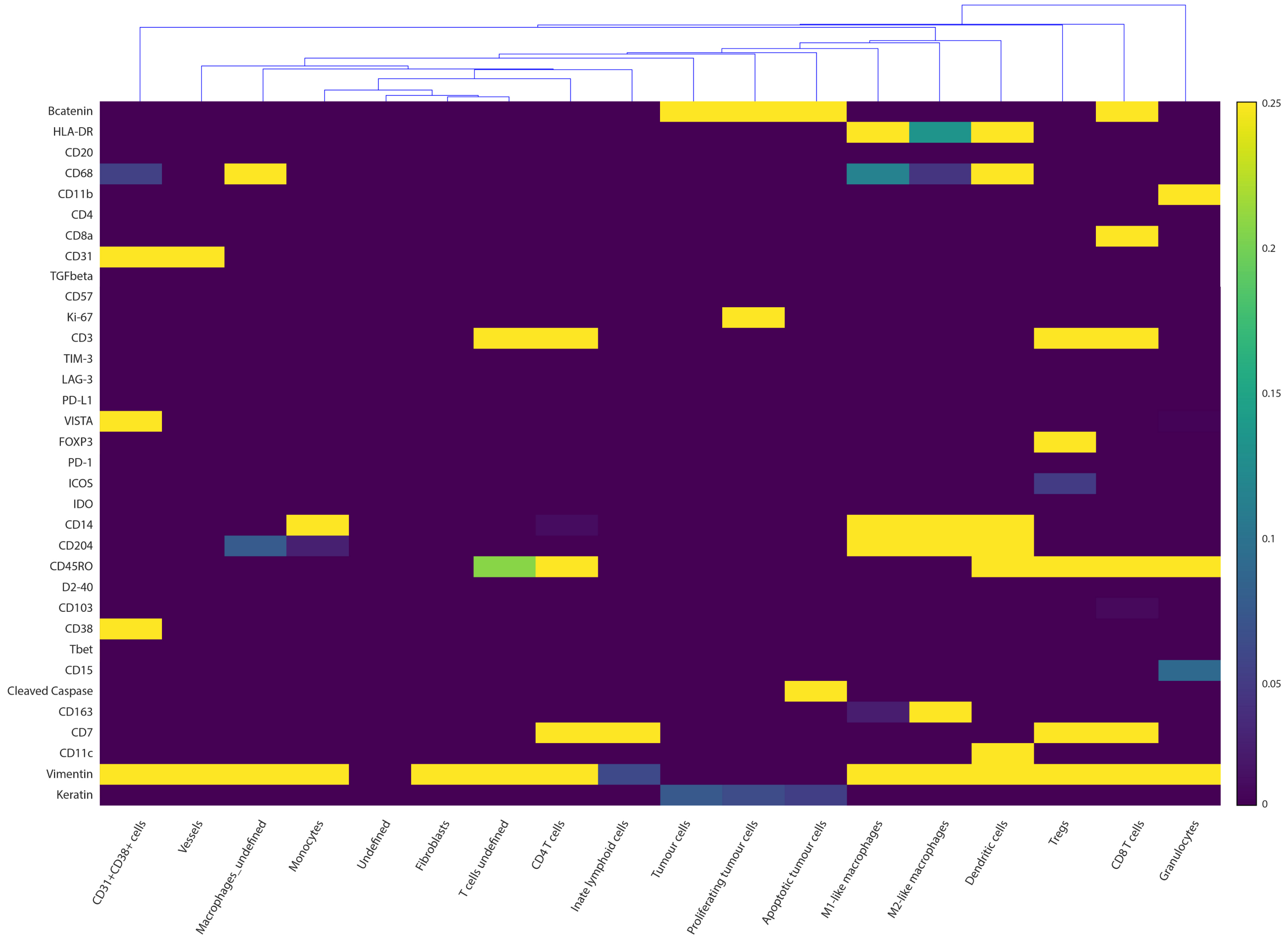


Supplementary Figure 5

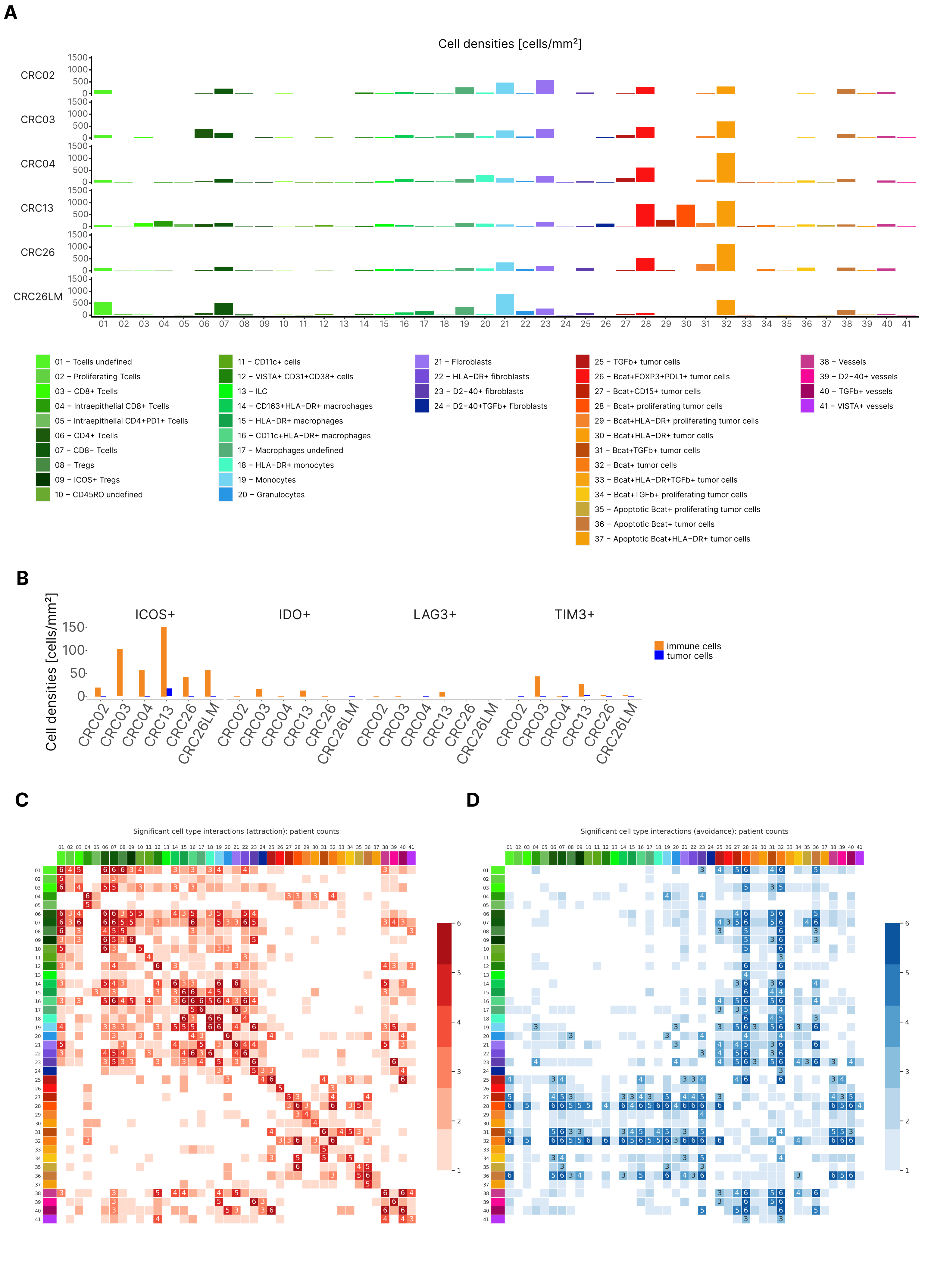


Supplementary Figure 6
